# MOFA-FLEX: A Factor Model Framework for Integrating Omics Data with Prior Knowledge

**DOI:** 10.1101/2025.11.03.686250

**Authors:** Arber Qoku, Martin Rohbeck, Florin C. Walter, Ilia Kats, Oliver Stegle, Florian Buettner

**Author notes:** These authors contributed equally (Arber Qoku, Martin Rohbeck, Florin C. Walter). These authors contributed equally (Oliver Stegle, Florian Buettner).

## Abstract

Latent factor models are first-line analysis approaches for single- and multi-omics data, essential for data integration, alignment, and biological signal discovery. To cater for new technologies and experimental designs, bespoke extensions of factor models have been proposed, incorporating spatial structure, temporal dynamics and the noise characteristics of single-cell assays. However, the development of tailored methods and software for individual use cases is laborious and requires advanced statistical and domain expertise, posing a significant barrier to users.

To address this, we here propose MOFA-FLEX, a flexible and modular factor analysis framework designed for customisable modelling across diverse multi-omics data scenarios. Built on probabilistic programming, MOFA-FLEX unifies previously isolated extensions of factor analysis – including flexible priors, non-negativity constraints, supervision signals, and alternative data likelihoods – allowing models to be configured declaratively without requiring manual engineering. Additionally, MOFA-FLEX features a novel domain knowledge module to inform and connect latent factors to gene programs.

We demonstrate MOFA-FLEX across multiple applications, showing (i) improved robustness in recovering gene programs from noisy prior knowledge in scRNA-seq data; (ii) effective disentanglement of technical and biological variation in multi-omic CITE-seq; and (iii) tailored spatial modelling that reveals spatially organised disease-associated gene programs in breast cancer.

## Introduction

The inference of interpretable molecular programs from high-dimensional omics data remains a central challenge ^1,2^. Advances in single-cell, multi-omics, and spatial profiling technologies have driven a surge in computational methods for integrating and interpreting different omics data ^3,4^. A widely used principle that underpins many of these methods is dimensionality reduction, whereby high-dimensional data are summarised in a lower-dimensional representation of latent factors or components ^5^. Although nonlinear deep learning models have recently gained traction ^6,7^, linear matrix factorization remains a widely-used first-line analysis approach owing to its inherent interpretability, directly revealing the mechanism by which latent factors explain the observed data variables. Additionally, factor analysis is particularly attractive for the analysis of typically smaller multi-omics patient cohorts, where highly parametrized deep-learning approaches struggle ^8^.

In recent years, numerous factor analysis models have been proposed, each extending the basic model in different directions, such as incorporating spatio-temporal dependencies ^9^, encouraging sparsity ^5,10^, enforcing non-negativity of the model weights and factors ^11,12^, incorporating supervision using external covariates ^13^, or accommodating prior knowledge to guide the inference of interpretable latent factors ^14–16^. While each of these methods are well-suited to their intended applications, transferring and combining the underlying modelling concepts across software tools is extremely challenging. The limited modularity of existing factor analysis frameworks constrains their adaptation to emerging data types and experimental designs. A key example is the challenge of modelling knowledge-guided factors while simultaneously capturing spatial or temporal structure in the data. Enabling rapid formulation of such combined models would open new opportunities to infer spatially localised pathway activity in tumour tissue or time-resolved immune response dynamics. A second example is the ability to combine structured sparsity with non-negativity constraints in the same model - two assumptions that have been individually considered and shown to be powerful for selecting biologically relevant features and producing interpretable additive factor representations. Critically, while these specific assumptions and many others have been considered in isolation in different specialised models, there is a lack of unifying framework that allows for combining these and other assumptions within the same model.

Here, we introduce MOFA-FLEX, a modelling framework that facilitates the rapid design of tailored matrix factorisation methods for a wide range of different multi-omics omics data types and designs - ranging from bulk to single-cell, and spatially resolved omics data. MOFA-FLEX generalises and unifies the capabilities of previous specialised factor models within a coherent framework and software. Instead of requiring separate tools for different tasks, users can declaratively compose models using a modular architecture that supports flexible priors, non-negativity constraints, supervised latent factors and alternative data likelihoods. This structure not only reproduces existing model classes as special cases but also extends them. Notably, MOFA-FLEX also introduces a new domain-knowledge module that allows informative gene programs to be embedded directly into the factor model via a structured sparsity prior. The modular design also allows for new components to be seamlessly incorporated, enabling future methodological extensions (e.g. alternative priors, supervision strategies, or data-driven constraints).

In summary, MOFA-FLEX provides a modular framework for factor analysis, for the first time allowing its users to encode rich modelling assumptions while preserving interpretability and enabling rapid adaptation to emerging omics technologies.

## Results

### MOFA-FLEX: A Modelling Framework for Multi-Omics Integration

MOFA-FLEX is a versatile Bayesian framework for multi-omics Integration via interpretable dimensionality reduction. The input data are structured multi-omics datasets in which observations (e.g., patients, samples, cells) – potentially belonging to multiple groups (e.g., different experimental conditions) – and variables (e.g., genes, genomic loci, metabolites) span multiple views (e.g., different omic modalities). For individual observation groups, additional observation-level covariates may be available, such as temporal or spatial information. Individual groups and views may exhibit distinct statistical properties, including count, binary, or continuous distributions, and may contain missing values - common in real-world datasets. MOFA-FLEX can account for gene set information associated with any of the input views, such as those from gene set annotation databases. A final factor analysis model is declaratively defined by the user, by choosing among existing modules, and how its outputs enable downstream biological interpretation (Fig. 1).

**Figure 1.**
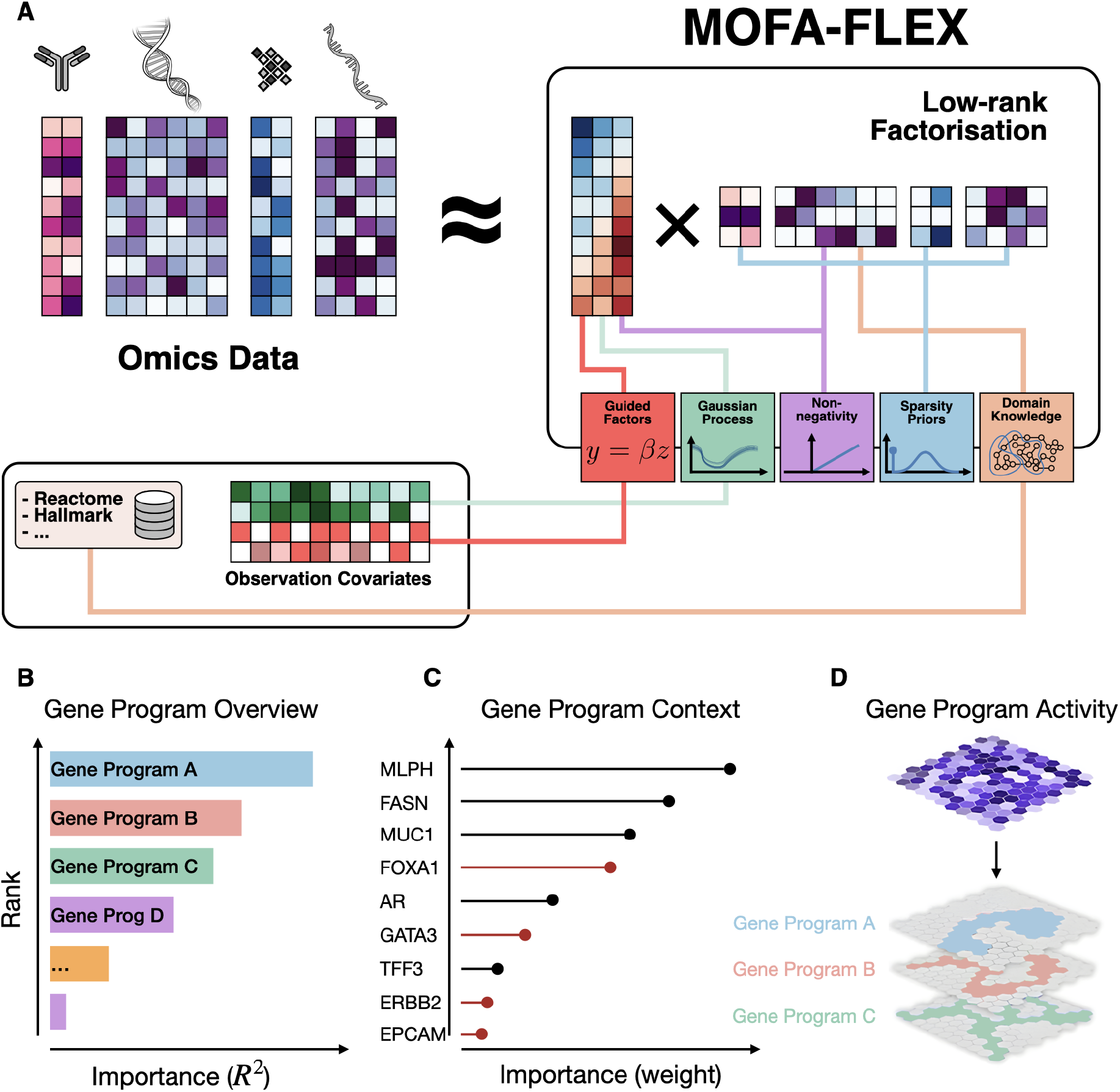
Overview of the MOFA-FLEX framework and downstream use cases. **(A)** Overview of the modelling approach. The MOFA-FLEX framework takes single or multi-omics data as input, alongside optional spatial or temporal covariates, supervision covariates, as well as domain knowledge about variables in the form of gene programs. The user can tailor models by flexibly combining modules (coloured boxes, right) for non-negativity, spatial covariates, covariate supervision and mechanisms to incorporate domain knowledge and sparsity assumptions. **(B-D)** A trained MOFA-FLEX model facilitates a number of downstream use cases, such as the investigation of the most relevant gene programs **(B)**, the contextualisation of existing gene programs with a refined set of genes **(C)** and the spatial distribution of the factor scores providing insights into regions of interest **(D)**.

The modularity of MOFA-FLEX allows the user to design models tailored to complex experimental designs. Users can define the number of data groups (or views) ^5,10,17,18^, define assumptions on sparsity or data non-negativity ^11^, use Gaussian Process priors for spatially or temporally resolved data ^9,12^, or employ supervised or guided factors ^13^ (Fig. 1A). MOFA-FLEX comes with ready-to-use implementations of modules derived from prior work (c.f. Methods; Supplementary Tab. S1). Owing to its modular design, these approaches can be flexibly combined and extended, thereby enabling novel combinations of assumptions. Additionally, MOFA-FLEX introduces a domain knowledge module, which provides a principled approach for integrating prior knowledge encoded as gene programs ^14^. MOFA-FLEX builds on Variational Bayesian inference, enabling estimation of the model evidence for a given dataset and thus principled model comparison and selection. Despite its flexibility, MOFA-FLEX is computationally efficient, outperforming existing methods in both runtime benchmarks and downstream analysis tasks (Fig. 2; Supplementary Fig. S2). A decision workflow and guidance for users to select modules and key parameters is provided in Methods (c.f. Methods; Supplementary Fig. S1).

**Figure 2.**
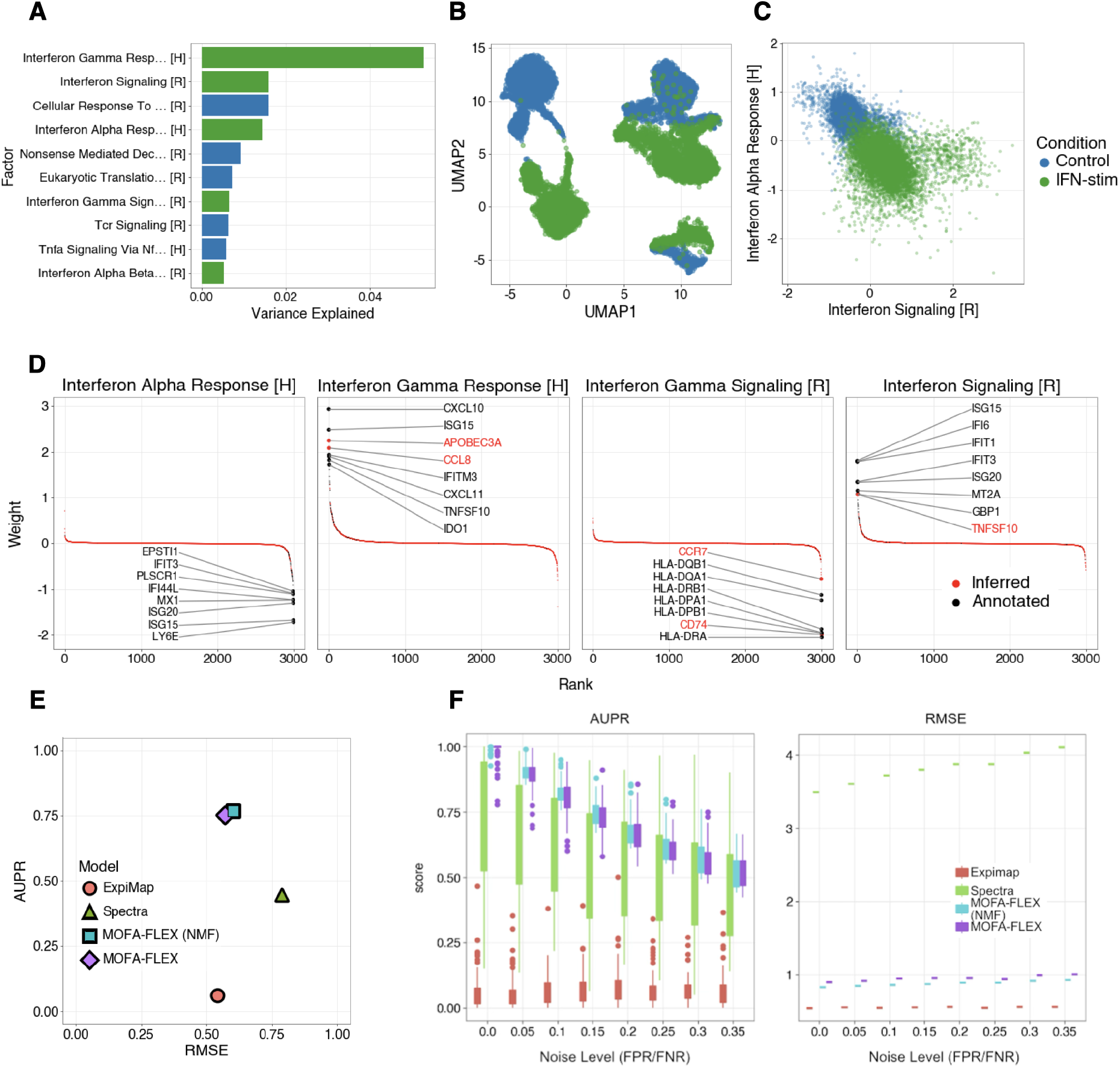
MOFA-FLEX identifies relevant gene programs and refines broad gene set annotations. Application of a single-view, single-group MOFA-FLEX configuration to a scRNA-seq dataset of peripheral blood mononuclear cells from lupus patients (N=13,576 cells). PBMCs were either treated with interferon-β (IFN-β) for ∼6 hours or left untreated. Domain knowledge was derived from publicly available gene set annotation resources, combining Hallmark and Reactome (65 gene sets total). **(A)** Inferred factors, ranked by the fraction of variance explained, labelled by their corresponding gene programs. Interferon-related programs highlighted in green. **(B)** UMAP representation of cells using the inferred latent factors as input; colour denotes stimulation status. **(C)** Scatter plot of factor activation for two representative IFN-related factors, labelled by their corresponding gene programs (cf. panel **A**), showing clear separation of IFN-stimulated vs control cells. **(D)** Gene weights for the top 8 genes of interferon-related gene programs. Genes that are part of the gene program are shown in black, and inferred genes in red. **(E)** Benchmark of gene program reconstruction based on semi-synthetic datasets derived from the same PBMC dataset. Gene set annotations from Hallmark and Reactome were deliberately perturbed by swapping 30% of genes in each set with genes outside of the set, thereby simulating noise that resembles gene-set misalignment in real gene set annotations. MOFA-FLEX, ExpiMap, Spectra, and MOFA-FLEX with nonnegative factors and weights, i.e., MOFA-FLEX (NMF) were considered for gene-program reconstruction. Shown is a scatter plot of the reconstruction RMSE, which quantifies data reconstruction performance, versus AUPR, which evaluates the reconstruction accuracy on the perturbed gene set annotations. **(F)** Analogous gene set reconstruction benchmark using synthetic data with known ground truth. Shown are AUPR (left) and RMSE (right) as a function of increasing levels of noise in the data generation process, randomly swapping up to 35% of genes in 5% increments.

A trained MOFA-FLEX model can be seamlessly used for different downstream analysis tasks (Fig. 1B-D). Users can identify the dominant gene programs driving inter-sample heterogeneity (Fig. 1B), explore the contextualisation of gene programs with novel genes (Fig. 1C), and visualise latent factor activity across spatial coordinates (Fig. 1D). The trained model can also be exported to R, enabling integration with existing R-based analysis pipelines, including those developed for the original Multi Omics Factor Analysis ^5^. Together, these capabilities enable MOFA-FLEX to serve both as an interpretability-first method and as a general-purpose modelling scaffold adaptable to diverse experimental designs.

### MOFA-FLEX Identifies Relevant Gene Programs and Reveals New Program-associated Genes

One of the primary motivations for employing factor analysis is their interpretability – either post-hoc by retrospectively annotating inferred factors ^5^, or by design using gene programs to directly capture biologically meaningful structure ^14–16^. As part of MOFA-FLEX, we introduce a novel domain knowledge module that connects inferred individual latent factors to predefined gene programs using a structured sparsity prior. To assess whether MOFA-FLEX can recover biologically meaningful gene programs, we considered a scRNA-seq dataset of peripheral blood mononuclear cells (PBMCs) ^19^ from lupus patients (N=13,576 cells), where cells were either treated with interferon-β (IFN-β) for ∼6 hours or left untreated. We configured MOFA-FLEX with a single group (cells) and view (RNA) To encourage the recovery of interpretable factors, we informed MOFA-FLEX with gene sets from the Reactome ^20^ and the Hallmark pathway database ^21,22^ via its domain knowledge module (c.f. Methods). Specifically, we ingested 52 gene sets from Reactome and 13 gene sets from the Hallmark collection. In addition, we included two uninformed latent factors to capture residual variation not explained by the informed factors. An overview of the MOFA-FLEX configuration across all applications is provided in Supplementary Table S2.

MOFA-FLEX prioritised gene programs consistent with the expected biology, ranking four interferon-related gene programs among the top factors (Fig. 2A). Additionally, the model recovered non-interferon programs known to be affected by IFN-β stimulation, such as Eukaryotic Translation Elongation ^23^. We compared MOFA-FLEX to existing methods for incorporating gene set information for interpreting single-cell RNA data: ExpiMap ^15^, a variational autoencoder that infers biological pathway-based latent components with a non-linear encoder and a linear decoder, and Spectra ^16^, a factorisation method that uses gene-gene knowledge graphs as biological context for optimisation. We applied ExpiMap and Spectra using the same gene sets, both of which notably did not prioritise interferon biology, despite IFN-related programs being present (Supplementary Fig. S4). A UMAP embedding of the MOFA-FLEX factors revealed a clear separation between IFN-stimulated and control cells (Fig. 2B). Notably, despite the added structure from prior knowledge, MOFA-FLEX retained the separation quality of an unconstrained PCA baseline, demonstrating that interpretability can be achieved without compromising expressive power (Supplementary Fig. S3). Inspection of individual gene program activations further confirmed that interferon-associated programs such as “Interferon Signaling” (Reactome) and “Interferon Alpha Response” (Hallmark) separated the experimental conditions (Fig. 2C).

Beyond prioritising gene programs that are consistent with known biology, the Bayesian modeling approach of the domain-knowledge module balances prior knowledge with the observed data, thereby allowing the refinement and the expansion of incomplete gene programs in a data-driven manner. MOFA-FLEX augmented the IFN programs by incorporating strongly supported genes absent from the curated sets (Fig. 2D). For example, the model recovered the interferon-stimulated gene TNFSF10 (TRAIL), which is induced by type I interferons to promote apoptosis in immune and non-immune cells as part of the antiviral and antitumour response ^24^. A comprehensive overview of the inferred gene programs a posteriori is provided in the supplementary materials (Supplementary Fig. S6).

The ability to refine gene programs improves the robustness to incomplete annotations—a common challenge for models relying on prior knowledge. We benchmarked MOFA-FLEX’s ability to recover original gene programs when given corrupted input sets and measured reconstruction error to assess model fit. This simulates real scenarios where only broad gene sets need refinement for specific experiments. To corrupt gene sets, we randomly replaced 30% of their genes with random, non-member genes. These benchmarks are limited to single-cell RNA-seq data and can’t address multi-view or covariate-rich datasets, which we explore later. Notably, while Spectra requires non-negative expression data, MOFA-FLEX flexibly includes this assumption, so we also tested a MOFA-FLEX model with non-negative constraints (MOFA-FLEX (NMF)).

To assess model performance in recovering the original gene programs, we extracted the latent factor weights for MOFA-FLEX and Spectra, and the linear decoder weights for ExpiMap. Recovery accuracy was quantified using the area under the precision-recall curve (AUPR), while reconstruction fidelity was measured by the root mean squared error (RMSE) between observed and reconstructed expression values. ExpiMap, MOFA-FLEX, and MOFA-FLEX (NMF) achieved comparable reconstruction performance (RMSE = 0.54-0.56), whereas Spectra performed substantially worse (RMSE = 0.78; Fig. 2E).

To further benchmark recovery against known ground truth, we generated semi-synthetic datasets derived from 10 Reactome and Hallmark gene programs under varying noise levels. Consistent with its robustness to noisy or incomplete priors, MOFA-FLEX outperformed both ExpiMap and Spectra across all noise regimes (Fig. 2F, Supplementary Fig. S5).

In summary, MOFA-FLEX accurately recovers gene programs from noisy annotations while maintaining high reconstruction fidelity, yielding biologically meaningful and context-aware representations.

### MOFA-FLEX Disentangles Unwanted Variation from Biological Signal in Multi-Omic CITE-seq Data

We next applied MOFA-FLEX to a CITE-seq dataset of immune cells from murine spleen and lymph nodes (N=14,870 cells). This dataset included both transcriptome (RNA) measurements and corresponding surface protein expression data (D=112 protein markers) in the same cells ^7^, collected in two independent batches on different days. The multi-modal dataset poses a realistic integration challenge due to substantial technical batch effects that may mask genuine biological signals. Our aim was to evaluate whether MOFA-FLEX could simultaneously integrate both modalities, isolate unwanted technical variation, and recover interpretable, cell type-specific gene programs.

To tackle this task, we configured a multi-view MOFA-FLEX model with domain knowledge and non-negativity modules (Supplementary Tab. S2). The RNA and protein modalities were treated as complementary views of the same cells, with biological domain knowledge in the form of gene sets provided exclusively to the RNA modality. In total, the model comprised 59 latent factors: 56 factors were informed by the Hallmark gene set collection from MSigDB ^20,21,25^ and cell type-specific gene programs from the Microwell-seq single-cell atlas ^26^. Three factors were left uninformed to serve as unconstrained components absorbing residual variation including batch-related variation not explained by the gene programs. This reflected our expectations that both inter-cell type differences and intra-cell type programs contribute to variation in these datasets.

A UMAP representation of the informed factors associated with gene programs indicated that these domain knowledge-guided factors faithfully captured biological heterogeneity induced by differences between cell types (Fig. 3A–B). In contrast, an analogous representation derived from the complete latent space – including both informed and uninformed factors – was subject to pronounced batch effects (Supplementary Fig. S7A), indicating that MOFA-FLEX can effectively leverage domain knowledge to aid the disentanglement of biological signals from technical variation. For comparison, we also considered MOFA-FLEX fit without domain knowledge, assuming uninformed sparse factors only as considered in other methods such as MOFA (Supplementary Fig. S7B). The resulting latent space of this model resulted in substantially lower disentanglement performance, which is consistent with reduced biological conservation (Supplementary Fig. S9) ^27^, indicating that the domain knowledge itself and not merely the assumption of sparsity leads to this improvement.

**Figure 3.**
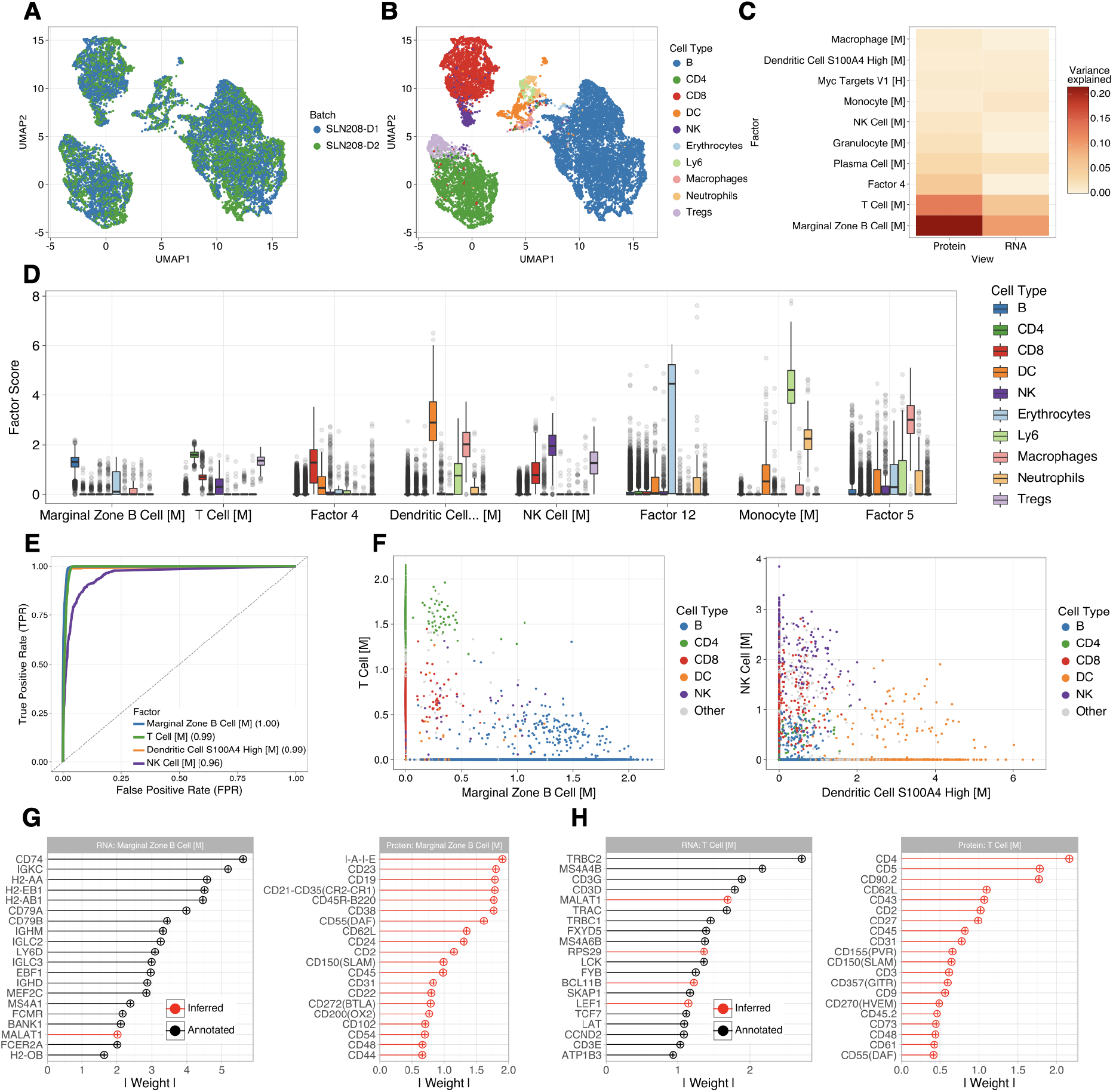
MOFA-FLEX disentangles batch and cell type variation from multi-omic CITE-seq data. Application of a multi-view MOFA-FLEX configuration to a multi-omic CITE-seq dataset from murine spleen and lymph nodes (N=14,870 cells), collected in two independent batches on different days. Domain knowledge was derived from publicly available gene set annotation resources, combining Hallmark and Microwell-seq single-cell atlas for mice (56 gene sets total). **(A, B)** UMAP based on the 21 most relevant gene programs, indicates control for batch effect **(A)** and preservation of cell type information **(B). (C)** Ranking of the inferred factors based on the variance explained, considering individual factors in each modality. As expected, factors informed by cell type gene programs a priori are the main drivers of variation. **(D)** Factor activity across 10 cell types, showing consistent activation of factors in the corresponding cell types. **(E)** Predictive capacity of four latent factors distinguishing their corresponding cell type from all others, quantified as area under the ROC curve (AUC). For instance, the latent factor informed by the “Marginal Zone B Cell” gene program clearly separates B cells. **(F)** Pairwise scatterplots of the activity of factors from **E. (G-H)** Top genes and surface proteins for the gene programs modelling B cells **(G)** and T cells **(H)**. Annotated genes (RNA level) are shown in black; inferred features in red. MOFA-FLEX refines both B cell and T cell gene programs and identifies the corresponding surface markers that are not covered in the prior annotation.

Overall, the model identified 21 active latent gene programs (retaining 99% of the explained variance) that captured the dominant sources of variation across modalities (Fig. 3C), providing a structured latent space suited for biological interpretation. At the same time, MOFA-FLEX preserved and amplified meaningful biological structure in an inherently interpretable manner. As expected, many of the most relevant factors directly corresponded to distinct immune cell populations, and hence were primarily active in the corresponding cell type, consistent with the cell-type gene programs used as priors (Fig. 3D). For example, the factor associated with the B cell gene program clearly delineated the B cells (Fig. 3E-F). Likewise, the factors associated with T cells, NK cells, and dendritic cells each showed high specificity for their respective cell identities. These results demonstrate that MOFA-FLEX can recover interpretable latent factors corresponding to known cell types, thereby providing a factorised and biologically meaningful representation of the cell-state landscape.

As part of the model inference MOFA-FLEX estimates the extent to which each factor is private to one view (active in one view only) or shared across views (active in RNA and protein view). The ability to model both views in this vein allows MOFA-FLEX to propagate knowledge from the informed RNA modality to the uninformed protein modality (Fig. 3G-H), which typically have too few markers to be informed directly. This configuration illustrates the utility of joint modelling across views and is not supported by existing single-modality gene program-based approaches that focus on scRNA-seq alone ^15,16^.

We again assessed the ability of MOFA-FLEX to refine and adapt broad gene program annotations to the specific context of this dataset, but now also included the protein layer that lacks prior information. The model preserved the global structure of the input gene programs, as confirmed by post hoc enrichment analysis using PCGSE ^28^, and simultaneously introduced additional genes. For instance, the marginal zone B cell gene program was expanded to include additional regulators such as PAX5 ^29^ and BCL11A ^30^ (Fig. 3G, Supplementary Fig. S8). Similarly, the T cell gene program was augmented with markers including BCL11B ^31^, LEF1 ^32^, IL7R ^33^, TXK ^34^, and NPC2 (Fig. 3H). The latent factor associated with CD4+ T cells identified CD4 as one of the top contributing genes among several T cell receptor components, demonstrating that MOFA-FLEX can reconfigure prior annotations in a meaningful manner (Supplementary Fig. S8). Complementary insights from the surface protein modality further validated these findings; shared latent factors highlighted canonical biomarkers such as the mouse I-A/I-E antibody and CD45R (B220) for B cells, while the T cell-associated factors prominently featured marker proteins including CD90.2, CD62L, CD5, and CD43.

Finally, to evaluate robustness to biologically misaligned priors, we provided a set of 55 gene programs related to spleen ageing from a different biological resource ^35^. MOFA-FLEX successfully adapted these inputs during training, reconfiguring them into the expected cell-type-specific programs characteristic of this dataset (Supplementary Fig. S10).

### MOFA-FLEX Maps Spatial Gene Program Activity in Breast Cancer

We next evaluated whether MOFA-FLEX could resolve spatially organised gene program activity in a complex tumour microenvironment. To this end, we jointly analysed two adjacent FFPE breast cancer tissue sections ^36^: one profiled with Chromium to obtain transcriptome-wide single-cell RNA-seq, and another profiled with Xenium to obtain spatially resolved expression of a targeted ∼300-gene panel. MOFA-FLEX was configured such that the broad transcriptomic coverage from Chromium informed gene program inference in Xenium, enabling spatial mapping of gene programs otherwise inaccessible due to limited panel size. This analysis illustrates how MOFA-FLEX can be flexibly tailored to complex designs, combining (i) a two-group architecture for sharing gene program weights across spatial (Xenium) and non-spatial (Chromium) views, (ii) a Gaussian Process prior to enforce spatial smoothness in Xenium, (iii) non-negativity constraints for interpretable factor loadings, and (iv) a gene set-informed prior to guide recovery of biologically meaningful programs. The configuration details for all applications are summarised in Supplementary Table S2.

The resulting model successfully inferred a rich latent representation that captured both global cell-type variation (Fig. 4A-B) and transcriptome-wide gene programs with spatially resolved activity (Fig. 4C-E; Supplementary Fig. S11). In particular, the Hallmark-informed factors revealed distinct spatial patterns corresponding to key breast cancer gene programs. Notably, the “Estrogen Response Early” (ERE) gene program – inferred using the larger transcriptomic coverage of the Chromium dataset – projected onto the Xenium spatial coordinates which cleanly separated invasive tumour regions from non-invasive areas. Cells with high ERE scores are predominantly localised within invasive niches, suggesting that elevated estrogen-driven transcriptional programs serve as indicators of tumour aggressiveness. Conversely, the “Apical Junction” (AJ) gene program primarily activated within ductal carcinoma in situ (DCIS 1 and DCIS 2) regions and was downregulated in invasive tissue, consistent with the loss of cell-cell adhesion during malignant progression. Beyond confirming expected patterns, MOFA-FLEX refined these annotations in a data-driven manner. Within the ERE module, classical markers such as TFF1 ^37,38^, GATA3 ^39^, and FOXA1 ^40^ emerged as top contributors, while genes associated with cell proliferation and metabolic reprogramming (e.g., FASN, CCND1, ERBB2) further delineated invasive phenotypes. Similarly, the AJ factor was enriched not only in established adhesion markers but also in novel candidates like CEACAM6 ^41^ and TACSTD2 ^42^, potentially offering novel insights into mechanisms governing tissue integrity and metastasis.

**Figure 4.**
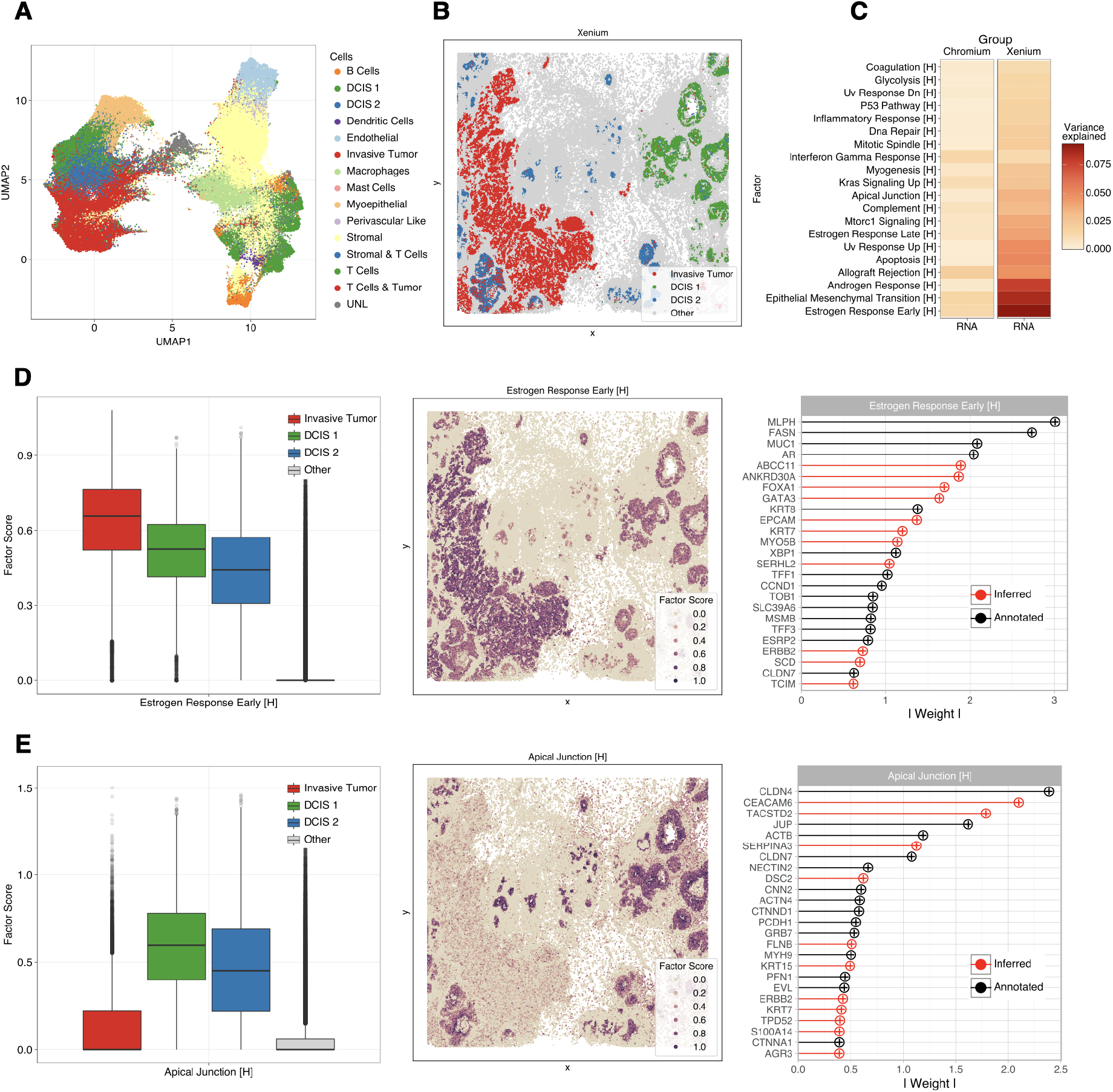
MOFA-FLEX delineates spatial gene program activity in human breast cancer. Application of MOFA-FLEX to breast cancer datasets comprising Xenium (166K cells; ∼300-gene spatial) and Chromium (30K cells; ∼3,000-gene non-spatial reference) data from adjacent FFPE slides. **(A)** UMAP of the cell types in the Xenium group. **(B)** Spatial mapping of cancer cell types, including invasive tumour and ductal carcinoma in situ (DCIS 1 and DCIS 2). **(C)** Most relevant latent gene programs ranked by variance explained across both groups. **(D)** “Estrogen Response Early” (ERE): factor scores across cancer cell types (left), spatial activity (middle), and top gene weights (right), with a-priori genes in black and inferred genes in red. MOFA-FLEX enriches ERE with known breast cancer markers such as FOXA1, GATA3 and ERBB2. **(E)** “Apical Junction” (AJ): factor scores (left), spatial activity (middle), and top gene weights (right). MOFA-FLEX enriches AJ with key genes such as CEACAM6 and TACSTD2, whose loss in invasive tumour cells promotes epithelial-to-mesenchymal transition (EMT).

In summary, MOFA-FLEX projects transcriptome-wide gene programs inferred from Chromium into spatial context via Xenium, enabling biologically interpretable, spatially resolved mapping of tumour organisation. This provides a high-resolution view of cancer architecture with direct relevance for mechanistic insight and translational application.

Finally, to illustrate its generality, we also applied MOFA-FLEX to additional datasets that were previously used to benchmark models such as MOFA ^5,10^, MEFISTO ^9^ and NSF ^12^. In each case, MOFA-FLEX faithfully reproduced the expected model behaviour and fully subsumed these methods, as their functionality can be recovered as special cases through the appropriate module configuration (c.f. Methods; Supplementary Fig. S12–S15). The implementation includes reproducible vignettes that demonstrate these configurations and serve as practical entry points for adapting MOFA-FLEX to new data and biological settings.

## Discussion

We present MOFA-FLEX, a new approach that elevates factor analysis from a set of specialised tools to a unified modelling framework for the omics community. Built upon a modular architecture grounded in probabilistic programming, MOFA-FLEX enables researchers to rapidly prototype and deploy custom factor models tailored to complex experimental designs and analysis tasks. This represents a fundamental shift in how latent variable modelling can be approached: instead of developing bespoke software for each new setting, users declaratively assemble models from a library of modules. At the same time, the framework is open to extension, allowing developers to contribute new modules as emerging analytical challenges arise.

Our case studies illustrate how this modularity translates into biological insight. In the breast cancer analysis, combining spatial smoothness, multi-group integration and non-negativity constraints revealed spatially organised gene program activity that would be inaccessible to monolithic tools. In the CITE-seq setting, shared factors enabled domain knowledge to propagate from the RNA modality into the protein space, achieving cross-modal interpretability without any custom algorithmic implementation. Each model was configured declaratively – no development of specialised architectures or custom algorithms was required.

Looking ahead, we envision MOFA-FLEX catalysing a community-driven ecosystem for modular factor analysis. Immediate opportunities include perturbation-aware models that disentangle treatment effects from technical confounders (e.g. CRISPR or drug response screens), modules that incorporate spatial adjacency graphs to capture tissue microenvironments, and hierarchical structures for multiscale biological organisation. The probabilistic foundation of the framework further provides a natural entry point for uncertainty quantification, offering principled ways to distinguish robust biological signal from technical noise in downstream predictive tasks.

In conclusion, MOFA-FLEX establishes a sustainable and forward-compatible framework for the community to advance multi-omics integration. As researchers contribute new modules for emerging data types and analytical objectives, the framework will evolve to meet future challenges while preserving the interpretability and statistical rigour that make factor models indispensable for biological discovery.

## Data Availability

A preprocessed version of the Kang dataset was obtained from [here](https://drive.google.com/uc?id=1t3oMuUfueUz_caLm5jmaEYjBxVNSsfxG).

A preprocessed version of the CITE-seq data was obtained from [here](https://github.com/YosefLab/totalVI_reproducibility/blob/master/data/spleen_lymph_111.h5ad) and [here](https://github.com/YosefLab/totalVI_reproducibility/blob/master/data/spleen_lymph_206.h5ad).

The CLL data was obtained from [here](https://ftp.ebi.ac.uk/pub/databases/mofa/cll_vignette/sample_metadata.txt) and [here](https://cf.10xgenomics.com/samples/spatial-exp/1.1.0/V1_Mouse_Brain_Sagittal_Anterior/V1_Mouse_Brain_Sagittal_Anterior_spatial.tar.gz).

The Mouse Brain Visium data was obtained from [here](https://cf.10xgenomics.com/samples/spatial-exp/1.1.0/V1_Mouse_Brain_Sagittal_Anterior/V1_Mouse_Brain_Sagittal_Anterior_filtered_feature_bc_matrix.h5) and [here](https://cf.10xgenomics.com/samples/spatial-exp/1.1.0/V1_Mouse_Brain_Sagittal_Anterior/V1_Mouse_Brain_Sagittal_Anterior_spatial.tar.gz).

The Mouse Brain Slide-seq v2 data was obtained from the SeuratData package.

Breast Cancer data was obtained from [here](https://doi.org/10.1038/s41467-023-43458-x) and [here](https://www.10xgenomics.com/products/xenium-in-situ/preview-dataset-human-breast).

## Code Availability

An open-source implementation of MOFA-FLEX is available on GitHub at https://github.com/bioFAM/mofaflex, where users can find detailed installation instructions. Additionally, comprehensive tutorials and documentation are available with supplementary materials for reproducing figures at https://github.com/bioFAM/mofaflex-analysis.

## Acknowledgements

This work has been supported by the European Research Council (Synergy Grant DECODE under grant agreement no. 810296), de.NBI (grant number W-de.NBI-013) and by the Deutsche Forschungsgemeinschaft (DFG, German Research Foundation) - 433034324.

This work was co-funded by the European Union (ERC, TAIPO, grant agreement No. 101088594 to F.B.). The views and opinions expressed are solely those of the authors and do not necessarily reflect those of the European Union or the European Research Council. Neither the European Union nor the granting authority can be held responsible for them.

## Author contributions

A.Q., F.C.W., M.R. and I.K. implemented the method, performed the analyses and generated the figures.

I.K. and A.Q. are the lead developers and maintainers of the package.

A.Q., F.C.W., M.R., F.B. and O.S. wrote the manuscript with input from I.K.

F.B. and O.S. conceived the project with input from A.Q., F.C.W. and M.R.

F.B. and O.S. supervised the project.

## Methods

Complete Methods are provided as supplementary file.

## Notes

https://github.com/bioFAM/mofaflex

### Competing Interest Statement

The authors have declared no competing interest.

