## Supplementary Information for "MOFA-FLEX: A Factor Model Framework for Integrating Omics Data with Prior Knowledge"

### Supplementary Figures

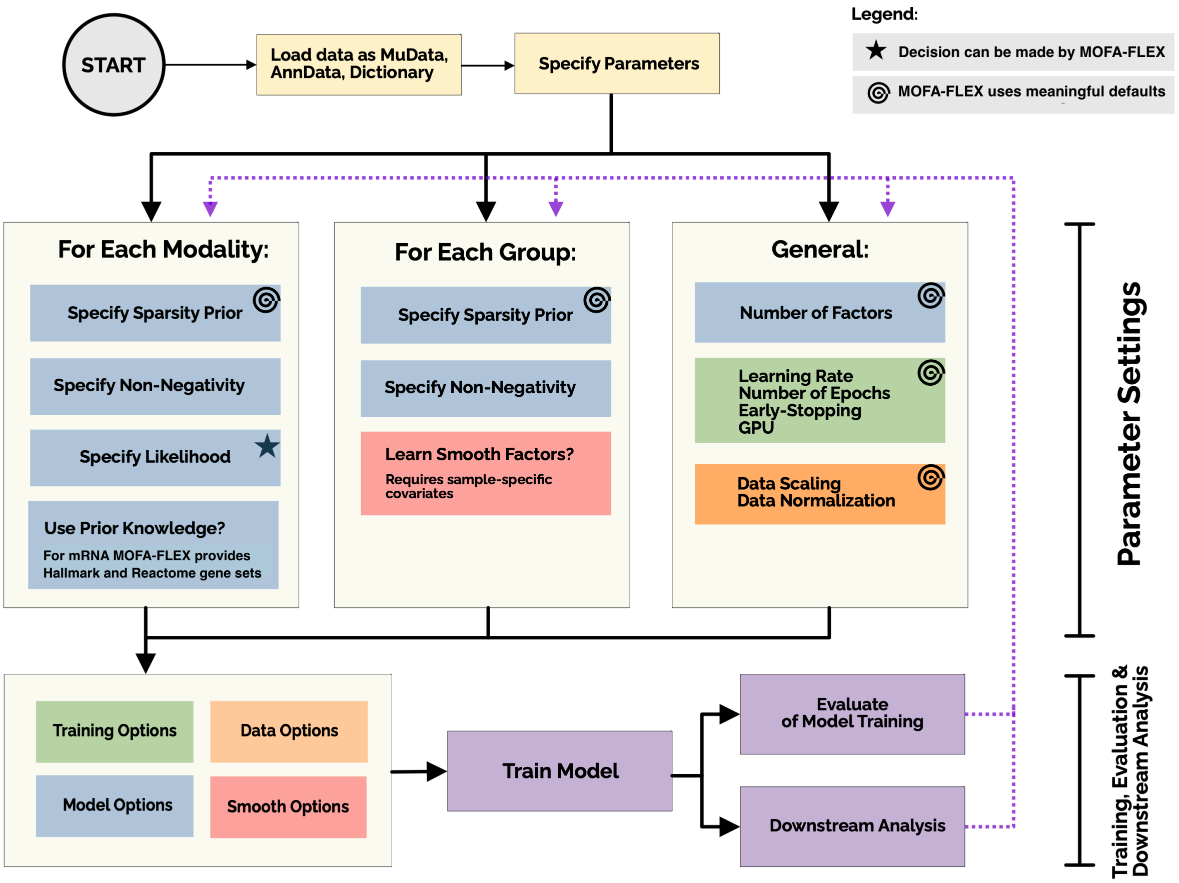

Figure S1: Decision workflow for configuring a MOFA-FLEX model. The input data are first loaded as MuData, AnnData or a standard Python dictionary. Users then define the model parameters in a modular manner. For each modality (e.g., RNA, ATAC, protein), a sparsity prior is selected from Laplace, (regularised) horseshoe or spike-and-slab; when informative gene programs are provided, the default is the regularised horseshoe. Non-negativity constraints may optionally be enabled to enforce positive feature weights or zero for improved interpretability. The data likelihood (Normal, Negative Binomial or Bernoulli) is automatically inferred during preprocessing but can be overridden if required. Next, prior biological knowledge can be specified — MOFA-FLEX provides access to the Hallmark and Reactome gene programs, but users may supply custom sets (e.g. via GMT files). For each group (e.g., patients, cells), users may independently configure sparsity priors and optionally enable spatial or temporal structure in the factor scores if such information is available. Finally, general configuration parameters are set, including the number of latent factors (beyond those informed by gene programs), training settings (learning rate, epochs, mini-batching) and data preprocessing options. Meaningful heuristic defaults are provided for all components, with options organised into clearly separated model, data, training (and optionally spatial/temporal) classes.

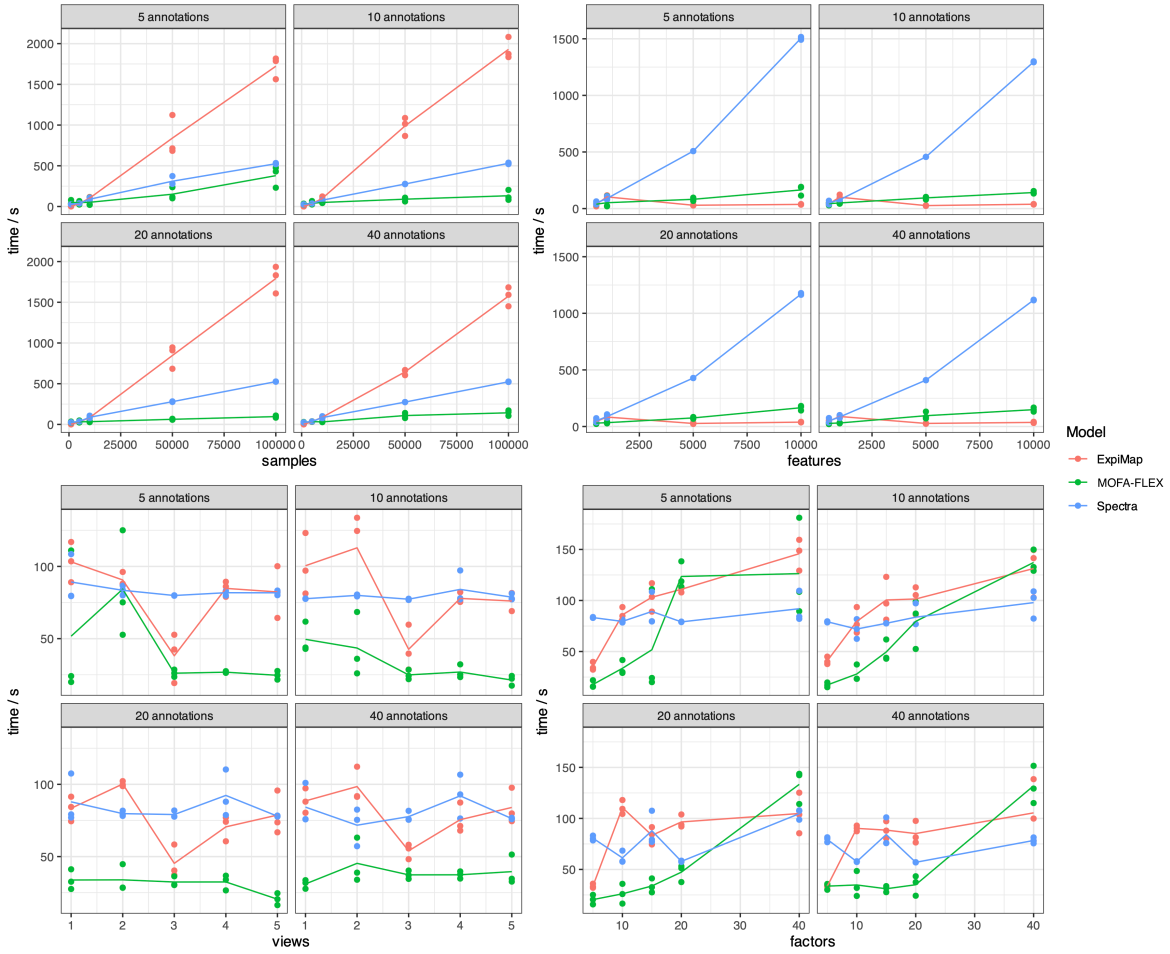

Figure S2: Runtime comparison between MOFA-FLEX, ExpiMap and Spectra. We benchmark all models on synthetic data across different annotations, sample sizes, factors, features and views. Time is measured in seconds and reported for three seeds. The synthetic data were generated using the MOFA-FLEX `DataGenerator` functionality. Briefly, factor and weight matrices were sampled from a Standard Normal distribution and multiplied. The result was used as the mean of a Normal distribution used for sampling the observations. To simulate real-world scenarios, we randomly turn off some latent factors in some views.

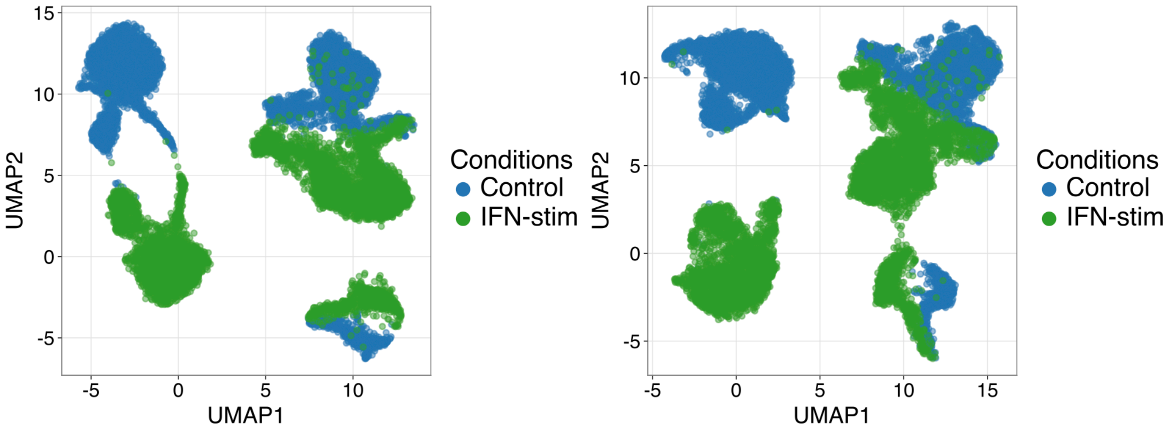

Figure S3: Decomposition comparison between MOFA-FLEX and PCA for the IFN-$\beta$ dataset [Kang et al., 2018]. Both methods yield a clear separation between IFN-stimulated and control cells in UMAP space. However, unlike the agnostic PCA baseline (right), MOFA-FLEX (left) achieves this separation via interpretable latent factors directly linked to biologically meaningful gene programs through its domain knowledge module.

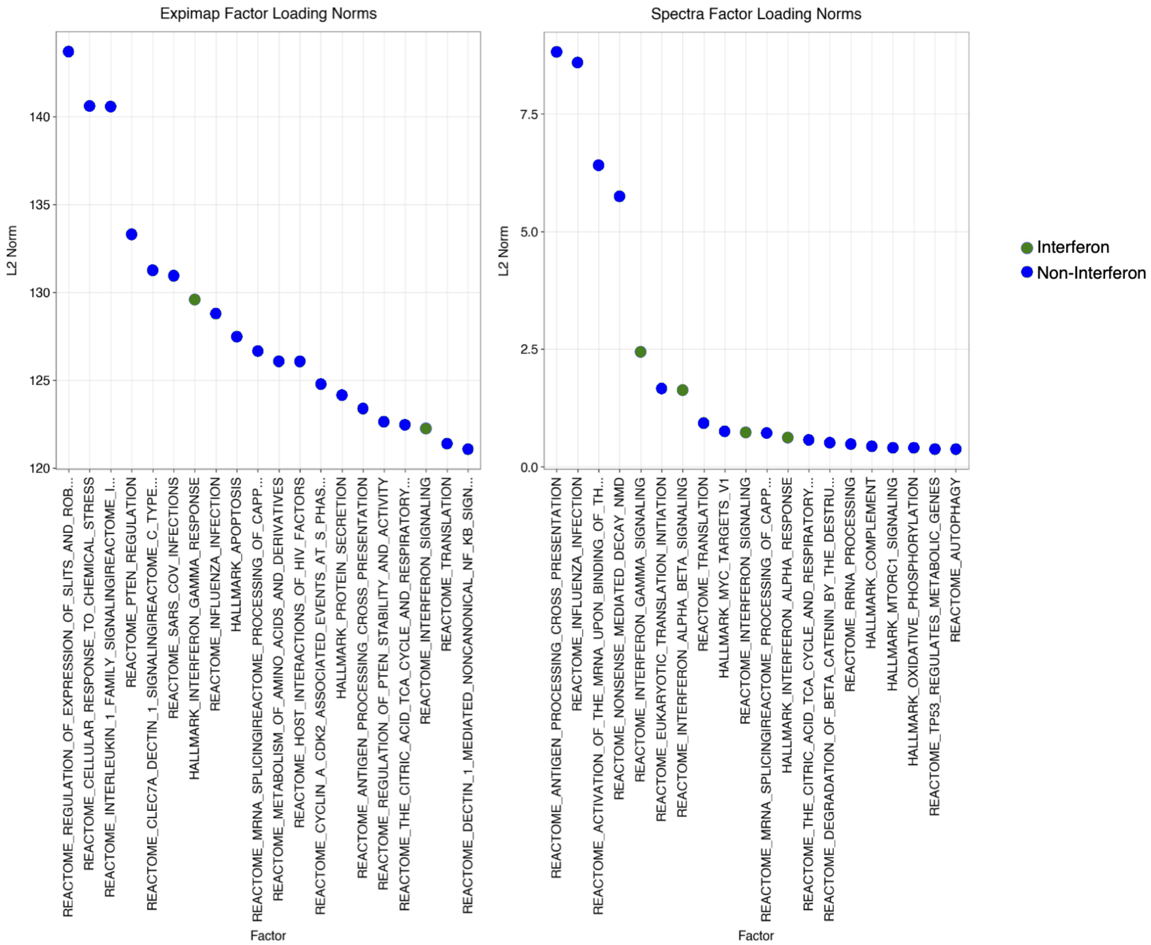

Figure S4: L2-Norms (y-axis) of factors (x-axis) for ExpiMap and Spectra (sorted by absolute value) for the IFN-$\beta$ [Kang et al., 2018] dataset. Interferon-named gene programs are coloured in green, and non-interferon-named gene programs are coloured in blue.

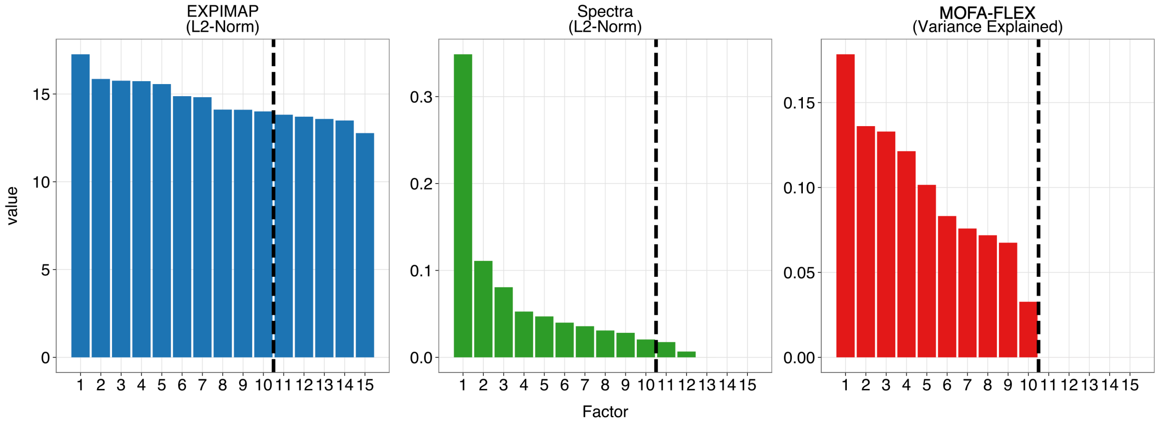

Figure S5: Sorted factor importance for three methods on synthetic data: ExpiMap, Spectra, and MOFA-FLEX. ExpiMap and Spectra factor importance is calculated using the L2-Norm, while MOFA-FLEX uses its variance explained computation. The dashed line indicates a potential cut-off for factor selection, given the ground truth number of factors is ten.

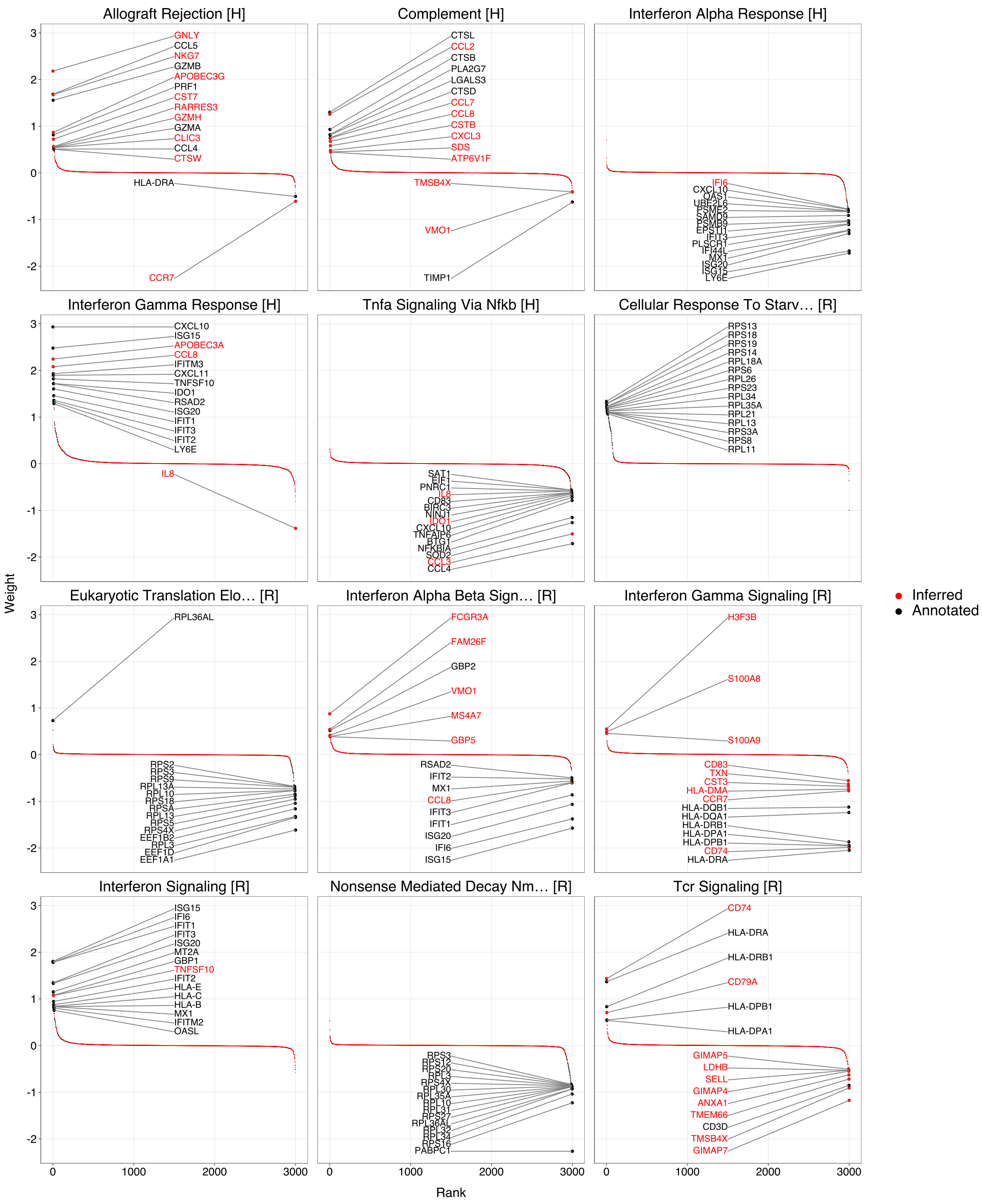

Figure S6: Hallmark [H] and Reactome [R] gene programs enriched with MOFA-FLEX for the IFN-$\beta$ [Kang et al., 2018] dataset. Genes coloured in red show the additional genes inferred with respect to the data context in each gene program among the top 12 gene programs with the highest rank according to variance explained.

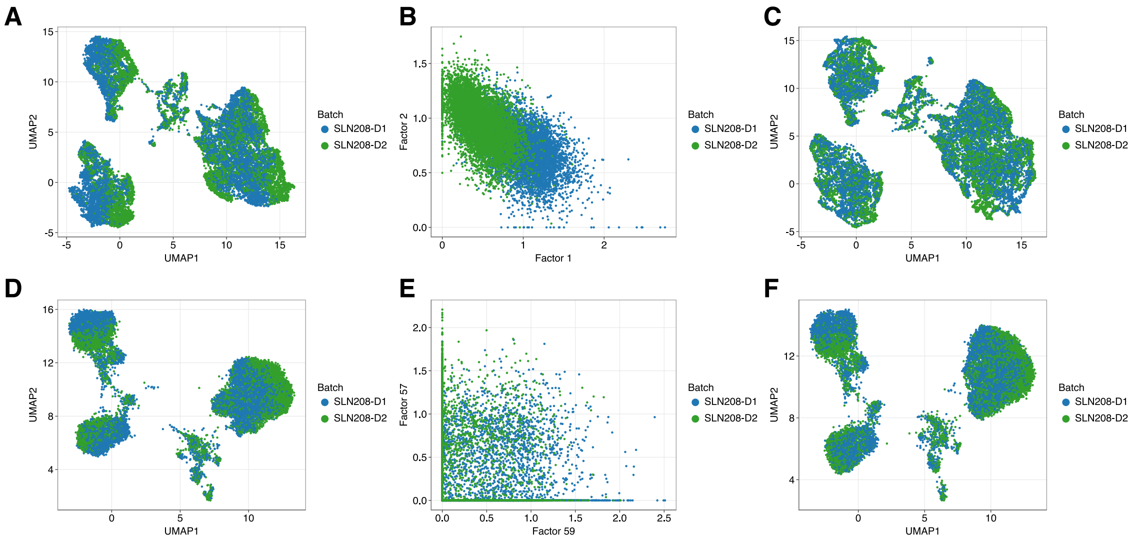

Figure S7: Comparison of MOFA-FLEX models on the CITE-seq murine data [Gayoso et al., 2021]. First row shows the results of the model trained on the informative mouse cell atlas (MCA) annotations, complemented by three uninformed factors as shown in the main paper. (**A**) The UMAP on the complete latent space — including the three uninformed latent factors — indicates the presence of the batch effect in the data. (**B**) The presence of the gene programs a priori renders the uninformed factors suitable for batch modelling. (**C**) Removing the three uninformed latent factors provides a better mixing of the batches. Second row shows the results of another MOFA-FLEX model trained with an uninformative prior (equivalent to MOFA). (**D**) The UMAP on the complete latent space indicates the presence of the batch effect in the data as in the previous model. (**E**) However, the uninformative prior is unable to model the technical variance from the batches within the first three latent factors. (**F**) Removing the top three most relevant factors for explaining the batch effect provides no improvement in mixing of the batches.

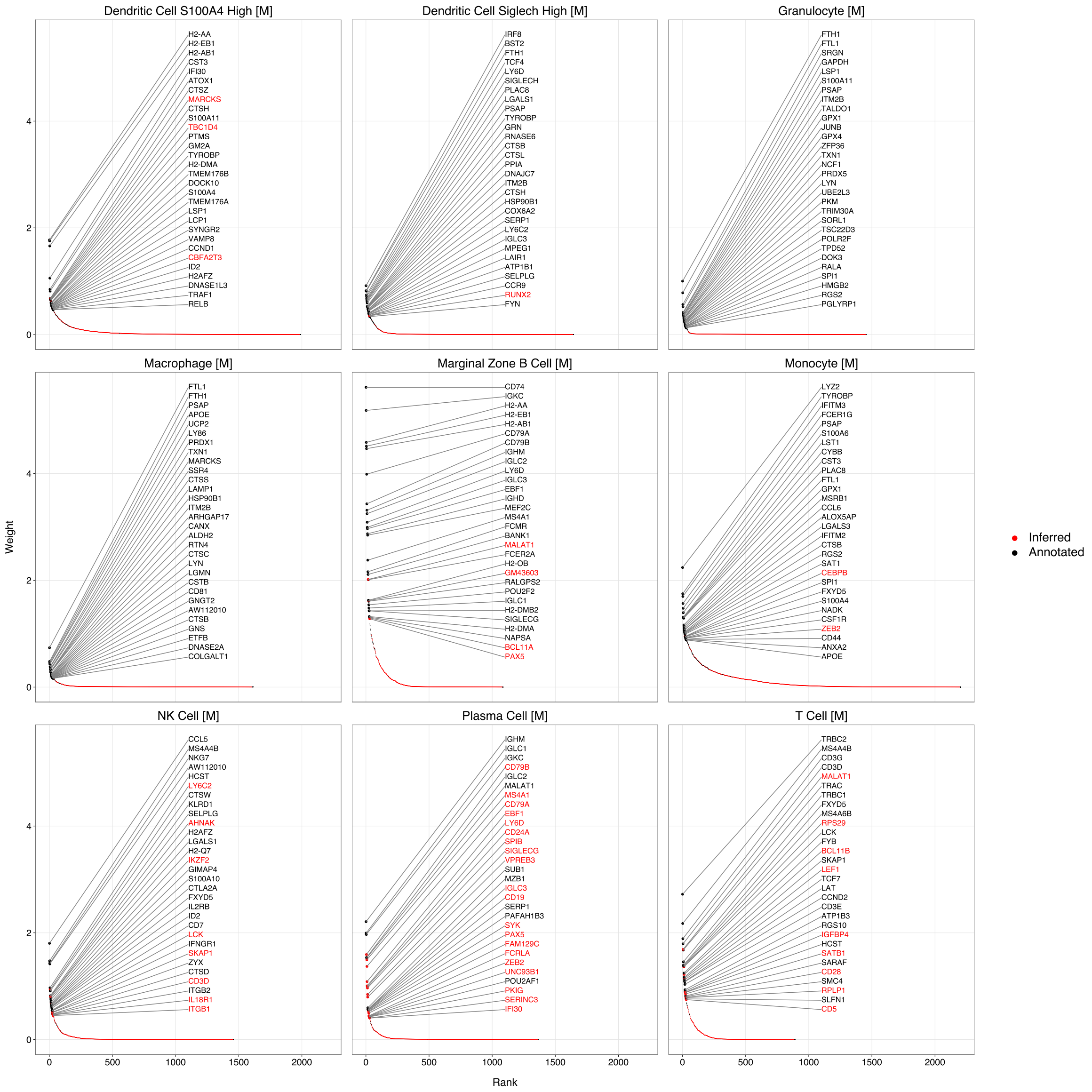

Figure S8: Cell-type gene programs enriched with MOFA-FLEX for the CITE-Seq [Gayoso et al., 2021] dataset. Genes coloured in red show the additional genes inferred with respect to the data context in each gene program among the top gene programs related to cell-type. The model preserved the global structure of the input gene programs, as confirmed by post hoc enrichment analysis using PCGSE, visible by the majority of the top genes being depicted in black. Simultaneously, MOFA-FLEX introduced additional genes with biological relevance to the corresponding gene programs. For instance, the marginal zone B cell gene program was expanded to include additional regulators such as PAX5 and BCL11A. Similarly, the T cell signature gene program was augmented with markers including BCL11B and LEF1.

| Method | Bio conservation | | | | | Batch correction | | | | | Aggregate score | | |
|---|---|---|---|---|---|---|---|---|---|---|---|---|---|
| | Isolated labels | KMeans NMI | KMeans ARI | Silhouette label | cLISI | Silhouette batch | iLISI | KBET | Graph connectivity | PCR comparison | Batch correction | Bio conservation | Total |
| **Informed [M]** | 0.57 | 0.79 | 0.71 | 0.64 | 1.00 | 0.90 | 0.61 | 0.60 | 0.87 | 0.97 | 0.79 | 0.74 | 0.76 |
| **Informed [T]** | 0.57 | 0.76 | 0.65 | 0.63 | 1.00 | 0.90 | 0.72 | 0.59 | 0.87 | 0.97 | 0.81 | 0.72 | 0.76 |
| **Uninformed** | 0.54 | 0.69 | 0.48 | 0.56 | 1.00 | 0.96 | 0.63 | 0.49 | 0.81 | 0.92 | 0.76 | 0.65 | 0.70 |
| **Unintegrated** | 0.57 | 0.58 | 0.37 | 0.64 | 1.00 | 0.77 | 0.00 | 0.27 | 0.85 | 0.61 | 0.50 | 0.63 | 0.58 |

Figure S9: Single-cell integration benchmarking (scIB) results for the CITE-Seq [Gayoso et al., 2021] dataset. We include two informed MOFA-FLEX models and compare to an uninformed MOFA-FLEX model (equivalent to MOFA), and an unintegrated PCA baseline. Informed [M] represents the MOFA-FLEX model shown in the main paper, informed by the combination of Hallmark and Microwell-seq Atlas [Han et al., 2018] gene programs. Informed [T] represents the MOFA-FLEX model informed by the combination of Hallmark and cell type gene programs specific to ageing processes [tab, 2020], as a noisy representation of the more suitable gene programs from the Microwell-seq Atlas. The MOFA-FLEX models equipped with domain knowledge, i.e. Informed [M] and Informed [T], achieve better results in both batch correction and biological conservation metrics when compare to the uninformed MOFA-FLEX model (equivalent to MOFA), and the unintegrated PCA baseline.

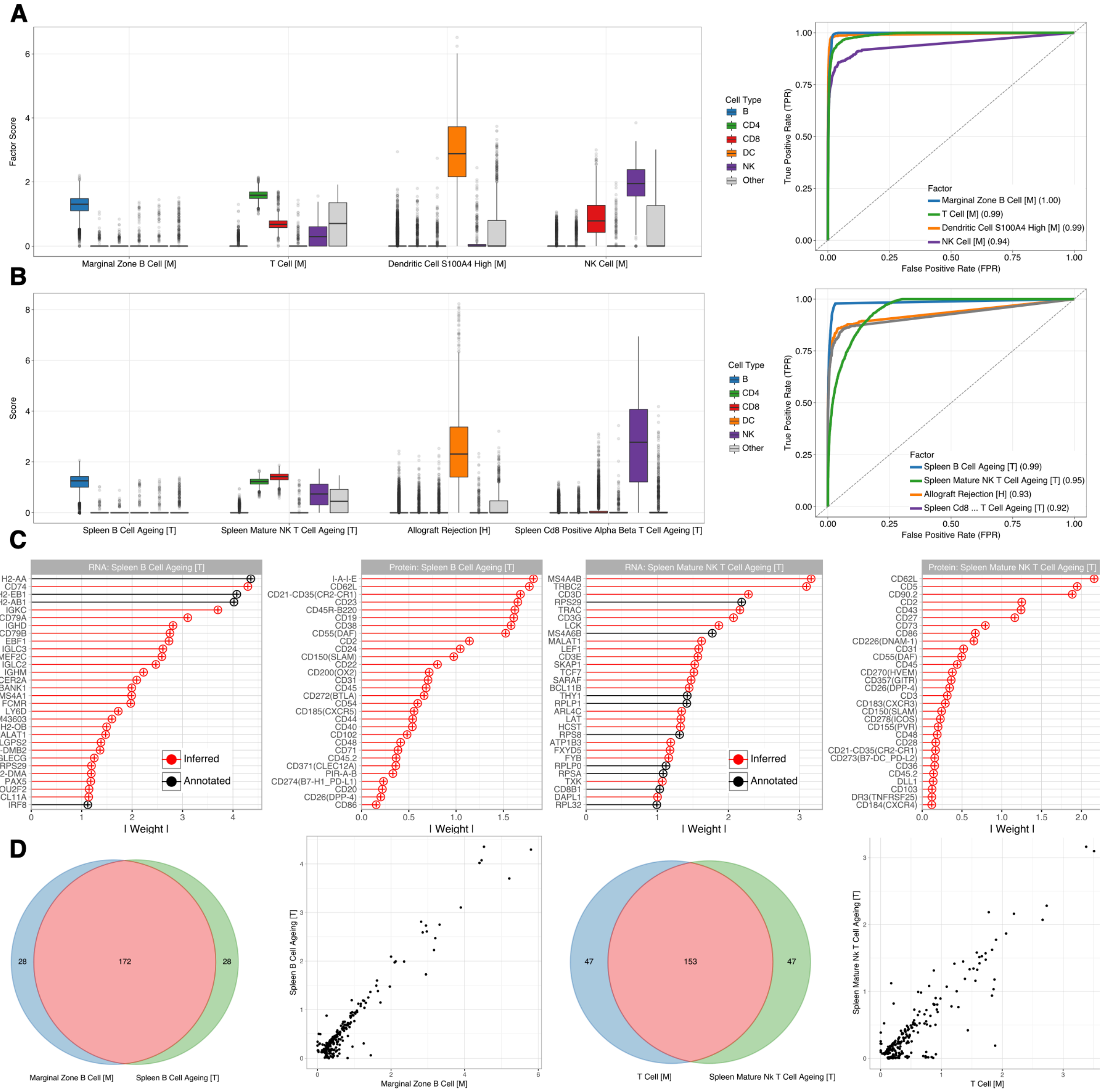

Figure S10: Comparison of different gene set priors for MOFA-FLEX trained on the CITE-seq data [Gayoso et al., 2021]. (**A**) First model ingesting the mouse cell atlas (MCA) as prior in the domain knowledge module (same as main paper); this provides suitable prior information as seen by the relevant gene programs differentiating their corresponding cell type (left), as quantified by the area under the ROC curve (right). (**B**) Second MOFA-FLEX model ingesting the organ-specific mouse gene set collection (Tabula Muris) into the domain knowledge module as a noisy version of the true underlying gene program. The corresponding scores given by the area under the ROC curve (right) suffer slightly in comparison to the MCA signature collection, but MOFA-FLEX is able to refine this noisy prior. (**C**) The top factor loadings for the gene programs modelling B cells and T cells. In particular, the B cell ageing gene program shows significant enrichment in the B cells and little to no enrichment for the rest of the cell population. Similarly, mature NK T cell ageing gene program is highly enriched in CD4 and CD8 cells. The factor loadings indicate major refinement of the prior gene set annotations, leading to difficulties in interpretation, unlike the more suitable MCA annotations which exhibit much less refinement. (**D**) The significant overlap between the top 200 factor loadings in both MOFA-FLEX models indicates that MOFA-FLEX is capable of converging to the true gene program composition regardless of the noise severity.

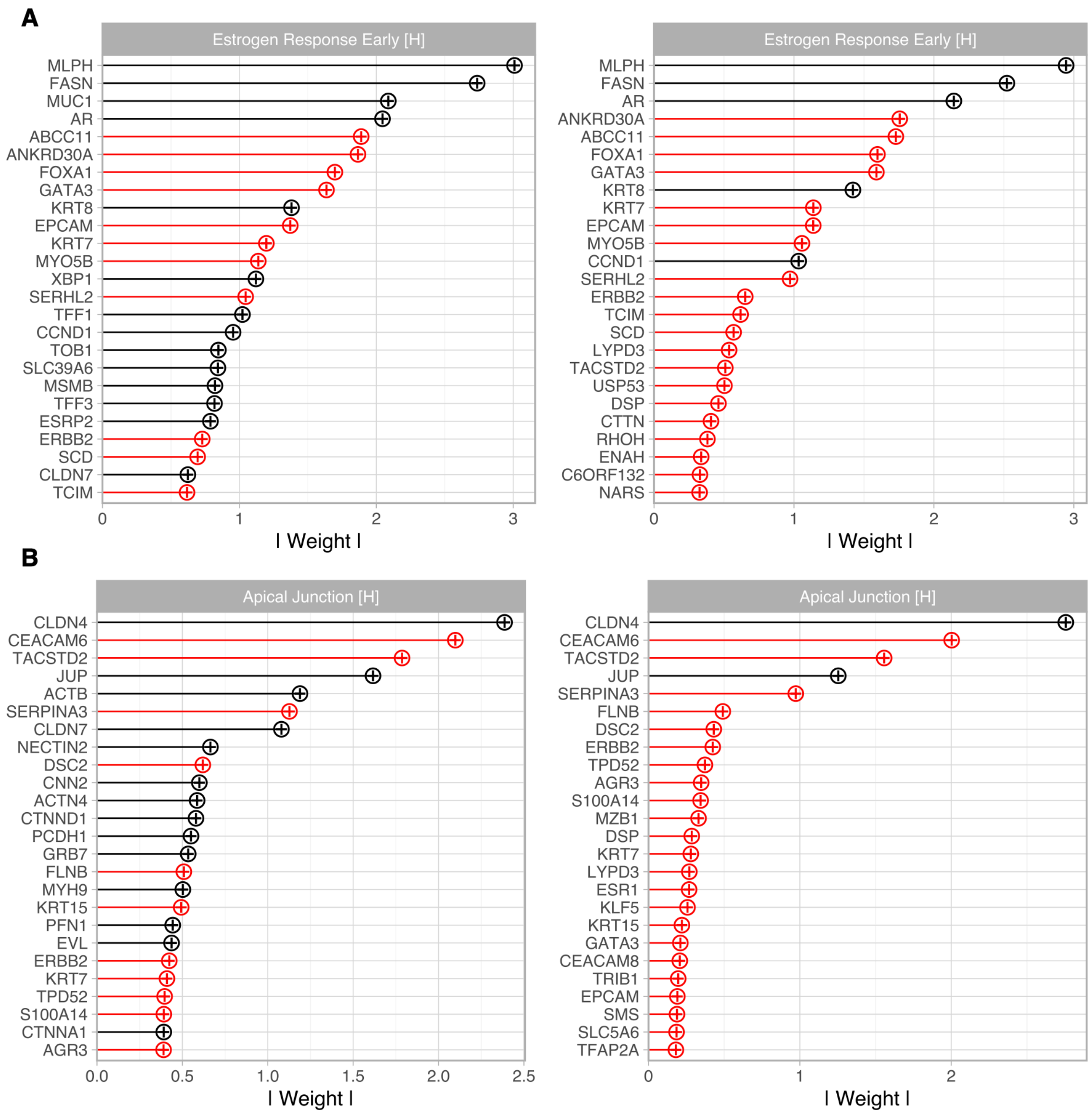

Figure S11: Comparison of the two main inferred gene programs in the breast cancer dataset [Janesick et al., 2023]. (A) The Estrogen Response Early gene program informed from the high coverage Chromium group leads to more robust adherence to the prior annotations (left), whereas having access to genes only available in Xenium (low coverage) leads to the majority of the loadings being inferred (right). (B) We observe similar results in the factor loadings a posteriori for the Apical Junction gene program. PCGSE renders the prior annotations accessing the Chromium gene space as significant for both gene programs.

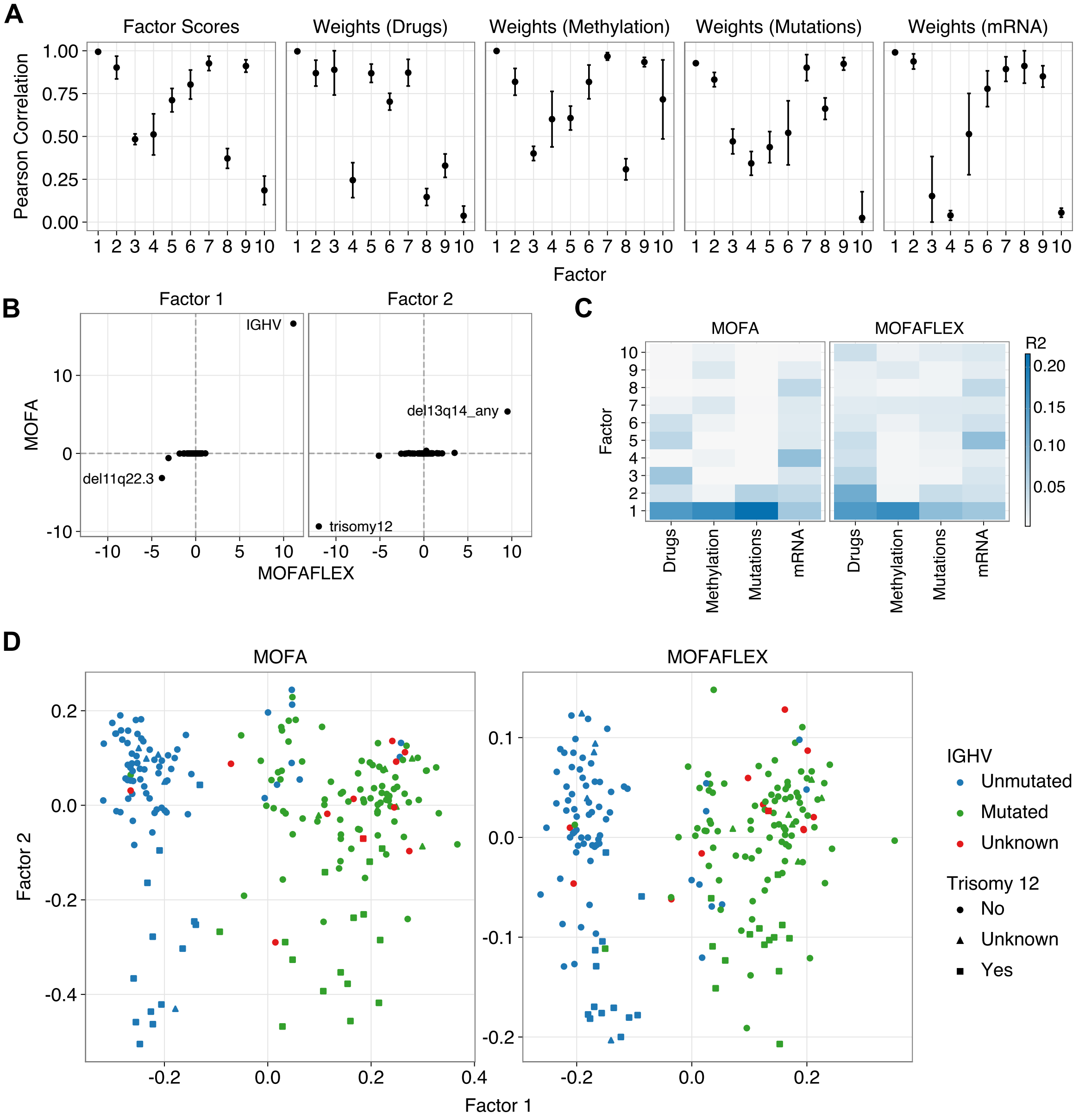

Figure S12: Application of MOFA and MOFA-FLEX to the CLL data. Both models were trained with 10 factors. (**A**) Pearson correlation coefficient of matched MOFA and MOFA-FLEX variables per matched factor. Errorbars show +/- one standard deviation from the mean based on 10 MOFA-FLEX random initialisations. In the following panels, one MOFA-FLEX initialisation is shown. (**B**) Inferred weights for the first two factors in the mutations view with MOFA-FLEX and MOFA values on the x and y axes, respectively. (**C**) Explained variance (R2, computed as in Argelaguet et al. [2018]) per factor and view in MOFA and MOFA-FLEX. (**D**) Factor 1 and 2 scores in MOFA and MOFA-FLEX, with colour and shape denoting IGHV and Trisomy 12 status, respectively.

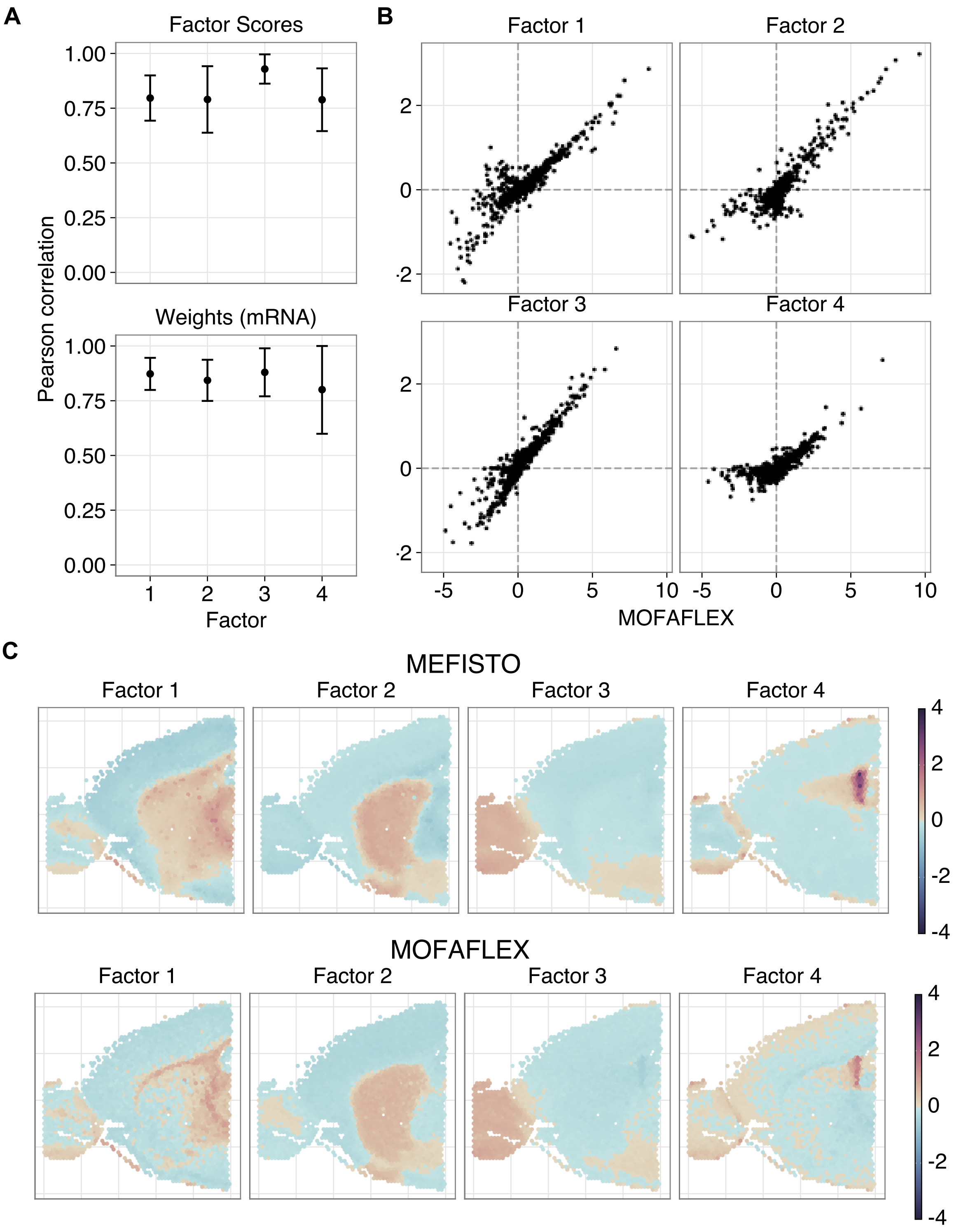

Figure S13: Application of MEFISTO and MOFA-FLEX to spatial transcriptomics data. Both models were trained with 4 factors. (**A**) Pearson correlation coefficient of MEFISTO and MOFA-FLEX latent variables per matched factor. Error bars show +/- one standard deviation from the mean based on 10 MOFA-FLEX random initialisations. In the following panels, one MOFA-FLEX initialisation is shown. (**B**) Scatter plots of MEFISTO and MOFA-FLEX weights per factor. (**C**) MEFISTO (top row) and MOFA-FLEX (bottom row) factor scores (colour) with spatial coordinates (x and y axes).

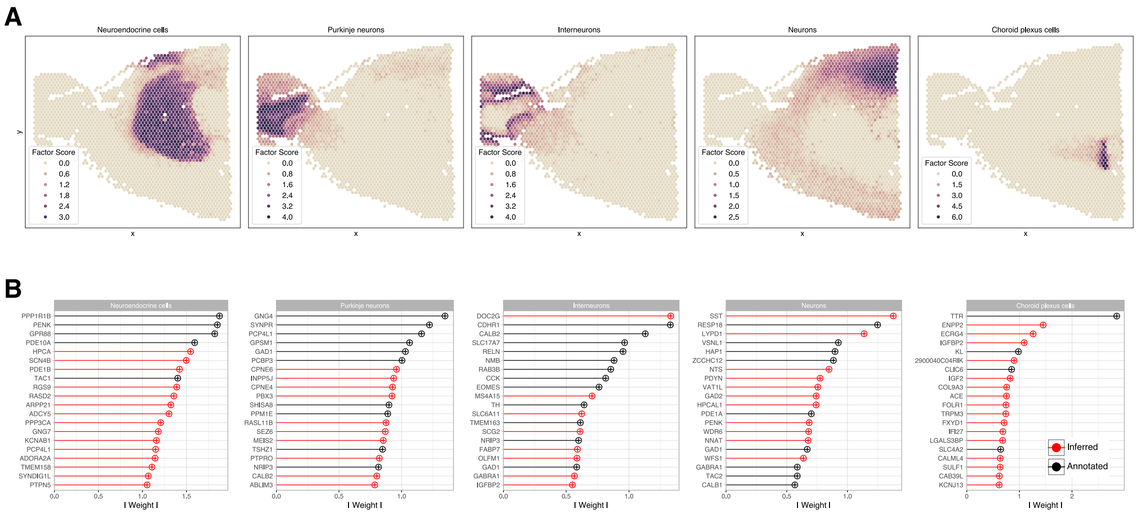

Figure S14: Application of MOFA-FLEX informed by brain cell type signatures from PanglaoDB [Franzén et al., 2019]. (**A**) Latent factor scores for the top factors informed by the cell type signatures a priori, mapped onto the spatial coordinates of the anterior mouse brain. (**B**) The corresponding factor loadings of each relevant latent factor ranking the top markers for each brain cell type. Markers in black depict the annotated features that were part of the signature a priori, whereas the markers in red depict the novel features that were added by the model during inference, thereby enriching the prior cell type signature with additional markers.

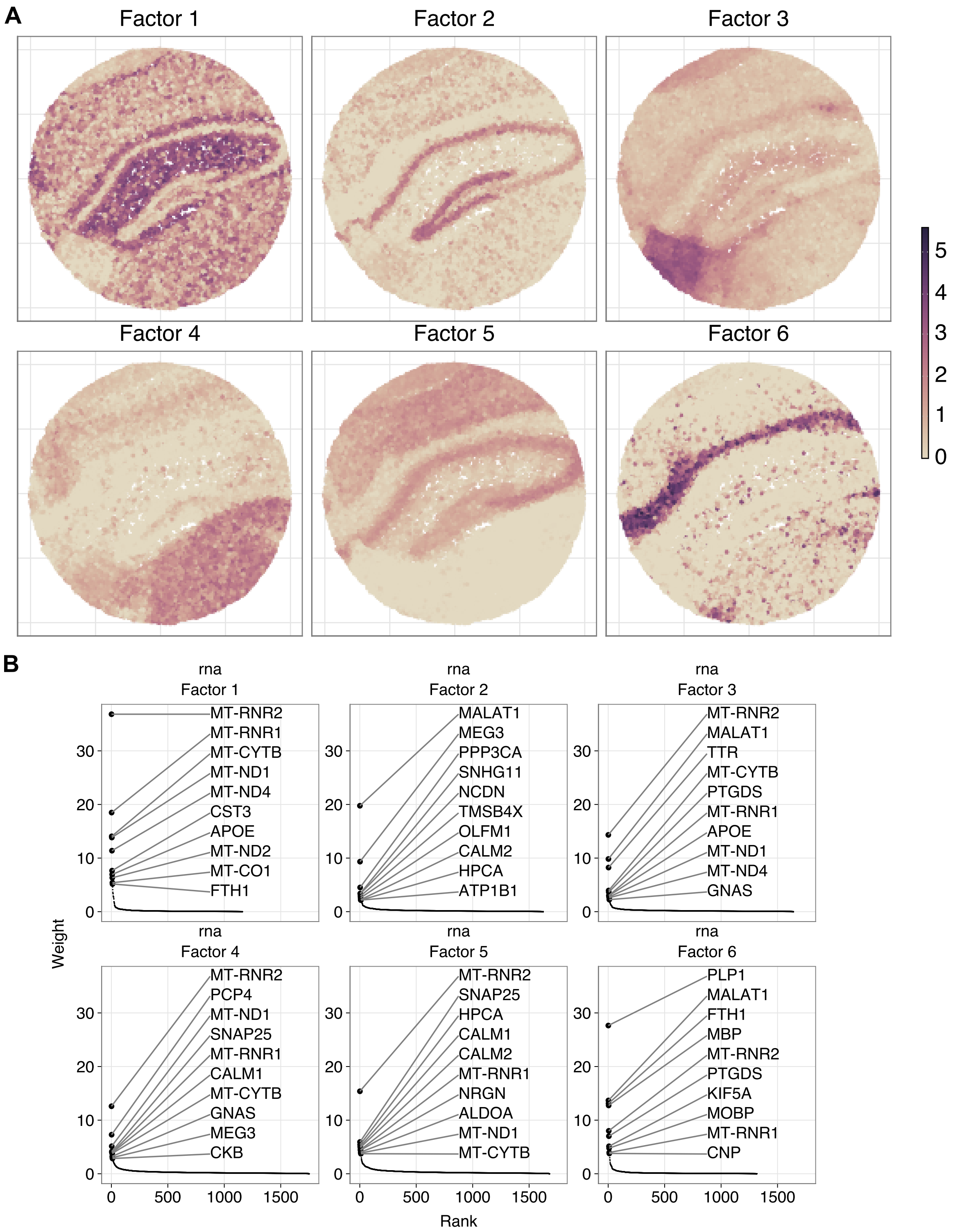

Figure S15: Application of MOFA-FLEX to the Slide-seq V2 data of the mouse hippocampus used in the NSF publication. The model was trained with 6 factors. (**A**) Factor scores (color) with spatial coordinates (x- and y-axes) and (**B**) factor weights.

### Supplementary Tables

Table S1: Comparison of features in different models. We compare ExpiMap [Lotfollahi et al., 2023], FISHFactor [Walter et al., 2023], F-scLVM [Buettner et al., 2017], GLM-PCA [Townes et al., 2019], MOFA [Argelaguet et al., 2018], MOFA+ [Argelaguet et al., 2020], MEFISTO [Velten et al., 2022], NSF [Townes and Engelhardt, 2023], SOFA [Capraz et al., 2024], OJSNMF [Esposito et al., 2019], Spectra [Kunes et al., 2024], ZIFA [Pierson and Yau, 2015] and ZINB-WAVE [Risso et al., 2018].

| Model | Sparsity | Domain Knowledge | Non-Negativity | Spatio-temporal | Multi-View | Multi-Group | Missing Observations | Guiding Variables |
|---|---|---|---|---|---|---|---|---|
| ExpiMap | $\ell_1$-norm | ✓ | | | | | | |
| FISHFactor | - | | ✓ | ✓ | | ✓ | | |
| f-scLVM | SnS | ✓ | | | | | ✓ | |
| GLM-PCA | - | | | | | | ✓ | |
| MOFA | SnS | | | | ✓ | | ✓ | |
| MOFA+ | SnS | | | | ✓ | ✓ | ✓ | |
| MEFISTO | SnS | | | ✓ | ✓ | ✓ | ✓ | |
| NSF | - | | ✓ | ✓ | | | ✓ | |
| SOFA | HS | | | | ✓ | | ✓ | ✓ |
| OJSNMF | $\ell_1$-norm | | | | ✓ | | | |
| Spectra | $\ell_1$-norm | ✓ | ✓ | | | | | |
| ZIFA | - | | | | | | | |
| ZINB-WaVE | - | | | | | | | |
| **MOFA-FLEX** | HS,SnS,Lap | ✓ | ✓ | ✓ | ✓ | ✓ | ✓ | ✓ |

Abbreviations: LAP: Laplace, SNS: Spike and Slab, HS: Horseshoe

Table S2: Module configurations used for each MOFA-FLEX application.

| **Dataset** | Sparsity | Domain Knowledge | Non-negativity | Spatio-temporal | Multi-view | Multi-group |
|---|---|---|---|---|---|---|
| IFN-$\beta$ [Kang et al., 2018] | HS | ✓ | | | | |
| CITE-Seq [Gayoso et al., 2021] | HS | ✓ | ✓ | | ✓ | |
| Breast Cancer [Janesick et al., 2023] | HS | ✓ | ✓ | ✓ | | ✓ |
| CLL [Dietrich et al., 2018] | HS | | | | ✓ | |
| 10X Visium [Stuart et al., 2019] | HS | | | ✓ | | |
| 10X Visium (Figure S14) | HS | ✓ | ✓ | ✓ | | |
| Slide-seq V2 [Satija et al., 2025] | HS | | ✓ | ✓ | | |

Table S3: Performance comparison of ExpiMap, MOFA-FLEX and Spectra. We report the average runtime per epoch and its standard deviation in seconds across three seeds.

| Factors | Samples | Features | Views | Annot. | ExpiMap | MOFA-FLEX | Spectra |
|---|---|---|---|---|---|---|---|
| 5 | 10 000 | 1000 | 1 | 5 | 35.34 ± 3.96 | 17.68 ± 3.60 | 83.25 ± 0.46 |
| 5 | 10 000 | 1000 | 1 | 10 | 40.85 ± 3.71 | 17.20 ± 2.33 | 79.15 ± 0.56 |
| 5 | 10 000 | 1000 | 1 | 20 | 33.76 ± 2.04 | 20.45 ± 4.69 | 80.62 ± 2.36 |
| 5 | 10 000 | 1000 | 1 | 40 | 33.51 ± 0.50 | 33.64 ± 3.11 | 79.42 ± 2.45 |
| 10 | 10 000 | 1000 | 1 | 5 | 85.95 ± 7.08 | 33.51 ± 7.05 | 79.48 ± 1.66 |
| 10 | 10 000 | 1000 | 1 | 10 | 79.48 ± 12.81 | 27.95 ± 8.10 | 72.03 ± 9.63 |
| 10 | 10 000 | 1000 | 1 | 20 | 110.57 ± 6.89 | 26.09 ± 9.69 | 61.32 ± 6.13 |
| 10 | 10 000 | 1000 | 1 | 40 | 90.24 ± 2.78 | 34.80 ± 12.47 | 57.68 ± 0.34 |
| 15 | 1000 | 1000 | 1 | 5 | 1.55 ± 0.12 | 51.49 ± 24.59 | 26.55 ± 0.19 |
| 15 | 1000 | 1000 | 1 | 10 | 1.56 ± 0.10 | 29.28 ± 6.04 | 25.47 ± 0.04 |
| 15 | 1000 | 1000 | 1 | 20 | 1.55 ± 0.15 | 26.76 ± 7.39 | 25.08 ± 0.04 |
| 15 | 1000 | 1000 | 1 | 40 | 1.64 ± 0.23 | 26.10 ± 3.46 | 24.55 ± 0.01 |
| 15 | 5000 | 1000 | 1 | 5 | 35.56 ± 3.53 | 47.85 ± 21.69 | 38.55 ± 0.10 |
| 15 | 5000 | 1000 | 1 | 10 | 32.99 ± 5.80 | 39.57 ± 24.05 | 36.98 ± 0.06 |
| 15 | 5000 | 1000 | 1 | 20 | 32.48 ± 6.30 | 32.71 ± 13.53 | 36.24 ± 0.07 |
| 15 | 5000 | 1000 | 1 | 40 | 34.06 ± 4.10 | 35.19 ± 3.28 | 35.59 ± 0.10 |
| 15 | 10 000 | 500 | 1 | 5 | 38.59 ± 17.10 | 39.11 ± 5.12 | 49.07 ± 13.26 |
| 15 | 10 000 | 500 | 1 | 10 | 41.79 ± 5.68 | 43.57 ± 13.52 | 50.29 ± 17.14 |
| 15 | 10 000 | 500 | 1 | 20 | 44.77 ± 9.66 | 29.97 ± 6.56 | 50.20 ± 20.15 |
| 15 | 10 000 | 500 | 1 | 40 | 47.34 ± 9.62 | 27.32 ± 6.55 | 48.89 ± 22.52 |
| 15 | 10 000 | 1000 | 1 | 5 | 103.17 ± 13.94 | 51.71 ± 51.43 | 89.21 ± 16.72 |
| 15 | 10 000 | 1000 | 1 | 10 | 100.49 ± 21.15 | 49.52 ± 10.63 | 77.64 ± 0.15 |
| 15 | 10 000 | 1000 | 1 | 20 | 83.42 ± 8.53 | 33.79 ± 6.89 | 87.90 ± 17.02 |
| 15 | 10 000 | 1000 | 1 | 40 | 88.54 ± 8.47 | 31.06 ± 3.04 | 84.24 ± 14.51 |
| 15 | 10 000 | 1000 | 2 | 5 | 90.61 ± 4.81 | 84.28 ± 37.03 | 83.48 ± 3.34 |
| 15 | 10 000 | 1000 | 2 | 10 | 112.87 ± 28.52 | 43.50 ± 22.20 | 79.97 ± 0.66 |
| 15 | 10 000 | 1000 | 2 | 20 | 100.24 ± 1.78 | 33.89 ± 9.39 | 79.73 ± 1.83 |
| 15 | 10 000 | 1000 | 2 | 40 | 98.48 ± 11.90 | 45.32 ± 15.60 | 71.71 ± 13.08 |
| 15 | 10 000 | 1000 | 3 | 5 | 38.15 ± 17.14 | 26.08 ± 2.50 | 79.87 ± 0.15 |
| 15 | 10 000 | 1000 | 3 | 10 | 42.64 ± 15.75 | 24.91 ± 3.33 | 77.30 ± 0.46 |
| 15 | 10 000 | 1000 | 3 | 20 | 45.31 ± 11.40 | 32.40 ± 3.43 | 79.10 ± 2.42 |
| 15 | 10 000 | 1000 | 3 | 40 | 54.00 ± 5.19 | 37.35 ± 2.90 | 77.66 ± 3.48 |
| 15 | 10 000 | 1000 | 4 | 5 | 84.80 ± 5.31 | 26.78 ± 0.62 | 81.80 ± 1.73 |
| 15 | 10 000 | 1000 | 4 | 10 | 77.87 ± 3.72 | 26.90 ± 4.68 | 84.10 ± 11.35 |
| 15 | 10 000 | 1000 | 4 | 20 | 70.56 ± 8.77 | 32.44 ± 5.25 | 92.33 ± 16.22 |
| 15 | 10 000 | 1000 | 4 | 40 | 75.53 ± 10.40 | 37.40 ± 2.49 | 92.04 ± 15.20 |
| 15 | 10 000 | 1000 | 5 | 5 | 82.31 ± 17.90 | 24.64 ± 2.99 | 81.73 ± 1.59 |
| 15 | 10 000 | 1000 | 5 | 10 | 76.04 ± 6.38 | 21.33 ± 3.47 | 78.73 ± 2.18 |
| 15 | 10 000 | 1000 | 5 | 20 | 78.74 ± 15.10 | 20.50 ± 4.12 | 77.96 ± 0.36 |
| 15 | 10 000 | 1000 | 5 | 40 | 84.00 ± 12.13 | 39.58 ± 10.29 | 76.31 ± 0.49 |
| 15 | 10 000 | 5000 | 1 | 5 | 29.40 ± 1.94 | 81.88 ± 14.52 | 507.76 ± 0.37 |
| 15 | 10 000 | 5000 | 1 | 10 | 26.41 ± 2.60 | 94.51 ± 13.19 | 456.65 ± 0.57 |
| 15 | 10 000 | 5000 | 1 | 20 | 26.81 ± 5.71 | 75.88 ± 8.06 | 428.31 ± 0.50 |
| 15 | 10 000 | 5000 | 1 | 40 | 27.16 ± 2.52 | 95.54 ± 31.92 | 409.45 ± 0.63 |
| 15 | 10 000 | 10 000 | 1 | 5 | 36.92 ± 2.54 | 164.75 ± 43.45 | 1505.32 ± 12.63 |
| 15 | 10 000 | 10 000 | 1 | 10 | 38.32 ± 2.12 | 141.85 ± 11.51 | 1295.62 ± 4.25 |
| 15 | 10 000 | 10 000 | 1 | 20 | 38.66 ± 2.53 | 165.93 ± 21.48 | 1169.32 ± 8.27 |
| 15 | 10 000 | 10 000 | 1 | 40 | 36.30 ± 5.13 | 148.88 ± 17.95 | 1117.51 ± 2.59 |
| 15 | 50 000 | 1000 | 1 | 5 | 840.10 ± 245.82 | 151.10 ± 74.34 | 310.44 ± 54.67 |
| 15 | 50 000 | 1000 | 1 | 10 | 989.49 ± 112.59 | 89.84 ± 24.76 | 276.10 ± 2.17 |
| 15 | 50 000 | 1000 | 1 | 20 | 846.42 ± 142.20 | 63.02 ± 6.08 | 279.40 ± 3.51 |
| 15 | 50 000 | 1000 | 1 | 40 | 645.20 ± 35.24 | 109.08 ± 31.48 | 274.35 ± 1.21 |
| 15 | 100 000 | 1000 | 1 | 5 | 1722.04 ± 138.65 | 378.91 ± 130.49 | 525.27 ± 7.54 |
| 15 | 100 000 | 1000 | 1 | 10 | 1929.84 ± 133.37 | 132.18 ± 63.75 | 530.54 ± 8.90 |
| 15 | 100 000 | 1000 | 1 | 20 | 1791.68 ± 166.81 | 96.19 ± 13.02 | 525.60 ± 1.26 |
| 15 | 100 000 | 1000 | 1 | 40 | 1575.22 ± 116.91 | 142.81 ± 34.07 | 523.74 ± 2.69 |
| 20 | 10 000 | 1000 | 1 | 5 | 110.96 ± 2.69 | 123.51 ± 13.14 | 79.03 ± 0.23 |
| 20 | 10 000 | 1000 | 1 | 10 | 101.46 ± 13.72 | 79.51 ± 24.12 | 83.63 ± 11.77 |
| 20 | 10 000 | 1000 | 1 | 20 | 96.55 ± 6.39 | 47.33 ± 8.48 | 57.90 ± 0.63 |
| 20 | 10 000 | 1000 | 1 | 40 | 85.15 ± 11.04 | 35.01 ± 9.78 | 56.99 ± 0.10 |
| 40 | 10 000 | 1000 | 1 | 5 | 145.84 ± 15.33 | 126.35 ± 48.33 | 91.92 ± 15.31 |
| 40 | 10 000 | 1000 | 1 | 10 | 131.30 ± 24.90 | 137.23 ± 11.11 | 97.92 ± 13.91 |
| 40 | 10 000 | 1000 | 1 | 20 | 104.79 ± 19.90 | 133.38 ± 16.70 | 104.63 ± 5.12 |
| 40 | 10 000 | 1000 | 1 | 40 | 105.44 ± 30.62 | 131.94 ± 18.41 | 78.37 ± 2.89 |

### Methods and Supplementary Information

Arber Qoku[1,4,8,*], Martin Rohbeck[3,7,*], Florin C. Walter[3,6,7,*], Ilia Kats[3], Oliver Stegle[3,6,7,#,†], and Florian Buettner[1,2,4,5,8,#,†]

[1]Institute of Informatics, Goethe University Frankfurt, Frankfurt am Main, Germany
[2]Department of Medicine, Goethe University Frankfurt, Frankfurt am Main, Germany
[3]Division of Computational Genomics and Systems Genetics, German Cancer Research Center (DKFZ), Heidelberg, Germany
[4]German Cancer Consortium (DKTK), partner site Frankfurt/Mainz, a partnership between DKFZ and UCT Frankfurt-Marburg, Germany, Frankfurt am Main, Germany
[5]Frankfurt Cancer Institute (FCI), Frankfurt am Main, Germany
[6]European Molecular Biology Laboratory (EMBL), Genome Biology Unit, Heidelberg, Germany
[7]Heidelberg University, Heidelberg, Germany
[8]Goethe University Frankfurt, Frankfurt am Main, Germany
[*]These authors contributed equally.
[#]These authors contributed equally.
[†]Corresponding authors.

**2 Existing Latent Variable Models** **30**

**3 Scalability Benchmarks** **30**

**4 Additional Results for PBMC Interferon Stimulated Dataset** **30**

**5 Handling Technical Variation and Noisy Priors in the Multi-omic CITE-seq Data 32**

**6 Overcoming Low Gene Coverage in Spatial Assays via Joint Modelling of Spatial and Single-cell Transcriptomics 32**

**7 Reproduction of MOFA, MEFISTO and NSF Results 34**

**8 Code Example 35**

### Mathematical Notation

| | |
|---|---|
| $x$ | A scalar value |
| $\mathbf{a}$ | A vector |
| $\mathbf{A}$ | A matrix |
| $\mathbf{A}^{\mathrm{T}}$ | Transpose of matrix $\mathbf{A}$ |
| $a_i$ | The $i$-th element of vector $\mathbf{a}$ |
| $A_{i,j}$ | The $i, j$-th element of matrix $\mathbf{A}$ |
| $\mathbf{A}_{i,:}$ | The $i$-th row of matrix $\mathbf{A}$ |
| $\mathbf{A}_{:,j}$ | The $j$-th column of matrix $\mathbf{A}$ |
| $\mathbf{1}_k$ | A vector of ones of length $k$. The subscript may be omitted if it is clear from the context |
| $\mathrm{diag}(\mathbf{x})$ | Diagonal matrix with the diagonal entries given by the vector $\mathbf{x}$ |
| $\mathbb{E}_q[X]$, $\mathbb{E}_{X\sim q}[X]$ | Expectation of the random variable $X$, where $X \sim q$. The subscript may be omitted if it is clear from the context |
| $\mathrm{KL}(p \parallel q)$ | Kullback-Leibler divergence of distributions $p$ and $q$ |
| $\mathcal{N}(\mu, \sigma^2)$ | Normal distribution with mean $\mu$ and variance $\sigma^2$ |
| $\mathrm{LogNormal}(\mu, \sigma^2)$ | Log-normal distribution with location $\mu$ and scale $\sigma$ |
| $C^+(l, s)$ | Half-Cauchy distribution with location $l$ and scale $s$ |
| $\mathcal{G}(\alpha, \beta)$ | Gamma distribution with shape $\alpha$ and rate $\beta$ |
| $\mathrm{Beta}(\alpha, \beta)$ | Beta distribution with parameters $\alpha$ and $\beta$ |
| $\mathrm{Ber}(p)$ | Bernoulli distribution with probability $p$ |
| $\mathrm{Cat}(p_1, \ldots, p_R)$ | Categorical distribution with category probabilities $p_r$ |
| $\mathrm{NB}(\mu, \alpha)$ | Negative binomial distribution with mean $\mu$ and overdispersion $\gamma$ |
| $\mathcal{GP}(m, v)$ | Gaussian process with mean $m$ and covariance $v$, where $m$ and $v$ can be functions |

### 1 Methods

MOFA-FLEX is a modular and unified Bayesian matrix factorization model designed for genomics applications. It integrates and extends previously published models to offer a comprehensive and user-friendly software package, emphasizing the incorporation of biological prior knowledge from databases to enhance inference and streamline downstream analyses.

As a flexible and modular framework, MOFA-FLEX is based on probabilistic matrix factorisation as detailed in Section 1.1.1. MOFA-FLEX can handle data from multiple data modalities and observation groups as described in Section 1.1.2 and provides multiple modules that users can combine freely to customize their analyses according to specific needs. For advanced users, MOFA-FLEX supports developing and integrating custom modules, enabling adaptation to emerging technologies and novel scientific questions. The currently pre-defined modules are:

- The *Sparsity Module* to obtain a sparse factorisation on the level of factors and individual variables (Section 1.2.1);
- The *Spatio-Temporal Module* to obtain spatially or temporally smooth factor scores (Section: 1.2.2);
- The *Non-negativity Module* to constrain factor scores and weight matrices to non-negative values (Section 1.2.3);
- The *Domain Knowledge Module* to inform the weight matrix with domain knowledge from data bases (Section: 1.2.4);
- The *Factor Supervision Module* to guide individual factors using external observation covariates (Section: 1.2.5).
- The *Likelihoods Module* to account for different data noise models (Section: 1.2.6).

#### 1.1 Factorisation Model

##### 1.1.1 Probabilistic Matrix Factorisation

MOFA-FLEX is based on Bayesian matrix factorisation, the probabilistic decomposition of a high dimensional data matrix $\mathbf{Y} \in \mathbb{R}^{N \times D}$ of $N$ observations and $D$ variables into the product of two lower rank latent matrices: The weight matrix $\mathbf{W} \in \mathbb{R}^{D \times K}$ and the factor scores matrix $\mathbf{Z} \in \mathbb{R}^{N \times K}$, where $K$ is the number of factors (or latent space dimensionality) and typically $K \ll D$. The objective is to infer a posterior distribution such that $\mathbf{Y} \approx \mathbf{Z}\mathbf{W}^T$, where $\mathbf{Z}$ and $\mathbf{W}$ represent point estimates from the posterior.
The Bayesian formulation of this model requires the specification of the latent variables' prior distributions and the choice of a likelihood function that probabilistically links the matrix product to the observed data (potentially including additional parameters). Assuming a general likelihood function $\mathcal{L}(\mu, \Theta)$ with mean parameter $\mu$ and a set of additional parameters $\Theta = \{\boldsymbol{\theta_1}, \boldsymbol{\theta_2}, \dots\}$, the factorisation of $\mathbf{Y}$ is given by

$$Y_{n,d} \sim \mathcal{L}(\mu = M_{n,d}, \Theta), \tag{1}$$

$$M_{n,d} = \sum_{k=1}^{K} Z_{n,k} W_{d,k}. \tag{2}$$

The versatility of models derived from this baseline matrix factorisation approach largely stems from the flexibility to impose structured priors $p(\mathbf{W})$ and $p(\mathbf{Z})$ or constraints on the latent matrices. This allows the posterior distribution to exhibit specific desirable properties, such as sparsity at the feature or factor level, covariance aligned with an external covariate, or alignment with prior knowledge from external databases.

##### 1.1.2 Multi-View and Multi-Group Structured Data

Multi-Omics Factor Analysis (MOFA/MOFA+) [Argelaguet et al., 2018, 2020] is a generalisation of Bayesian matrix factorisation to $V \times G$ data matrices $\{\mathbf{Y}^{v,g}\}_{v=1,\dots,V;g=1,\dots,G}$, where the index $v$ represents data *views* (or *modalities*) with different sets of features but at least partially overlapping observations and the index $g$ represents data *groups* with different sets of observations but at least partially overlapping features. Data views can be different omics profiling techniques applied to the same samples, e.g., transcriptomics, proteomics, or epigenomics, while data groups can be samples that were collected from different donors. The joint factorisation of the set of matrices results in view-specific weight matrices $\{\mathbf{W}^v\}_{v=1,\dots,V}$ that are shared between groups, and group-specific factor scores matrices $\{\mathbf{Z}^g\}_{g=1,\dots,G}$ that are shared between views. The factorisation of $\mathbf{Y}^{v,g}$ in this general case of multiple views and groups is given by

$$Y^{v,g}_{n,d} \sim \mathcal{L}(\mu = M^{v,g}_{n,d}, \Theta), \tag{3}$$

$$M^{v,g}_{n,d} = \sum_{k=1}^{K} Z^g_{n,k} W^v_{d,k}. \tag{4}$$

For ease of notation, the view and group indices $v$ and $g$ are in the following sections omitted for improved readability.

#### 1.2 Modules

In this section, we describe all currently available MOFA-FLEX modules that users can freely combine to tailor the analysis to their specific needs.

##### 1.2.1 Sparsity Module

To fully leverage the multi-view and multi-group structure, MOFA implements structured sparsity prior distributions for the weights and factor scores matrices. These priors encourage a high proportion of zero-valued entries on (a) the level of individual variables or observations and (b) the level of factors in different views or groups.

**Spike-and-Slab and Automatic Relevance Determination Prior** In MOFA, a combination of the Automatic Relevance Determination (ARD) prior [MacKay, 1996] for factor level sparsity with a Spike-and-Slab (SNS) prior [Mitchell and Beauchamp, 1988] for variable or observation level sparsity is used for this purpose. For a general latent matrix element $A_{i,j}$, this hierarchical prior is defined as

$$A_{i,j} = \hat{A}_{i,j} S_{i,j}, \tag{5}$$

$$\hat{A}_{i,j} \sim \mathcal{N}\big(0, 1/\alpha_j\big), \tag{6}$$

$$S_{i,j} \sim \mathrm{Ber}\big(\theta_j\big), \tag{7}$$

$$\theta_j \sim \mathrm{Beta}\Big(a_0^\theta, b_0^\theta\Big), \tag{8}$$

$$\alpha_j \sim \mathcal{G}(a_0^\alpha, b_0^\alpha), \tag{9}$$

where $a_0^\theta = b_0^\theta = 1$ and $a_0^\alpha = b_0^\alpha = 10^{-3}$ are constants with these default values in MOFA-FLEX. The weights and factor scores priors are special cases of this general prior, with $A_{i,j} = W_{d,k}$ for the weights or $A_{i,j} = Z_{n,k}$ for the factor scores. For multiple views or groups, a separate prior with the corresponding latent variables is used for each view and group. For more details, the reader is referred to the supplementary methods of the original publications [Argelaguet et al., 2018, 2020].

**Horseshoe Prior** A difficulty with the formulation of a sparsity prior in Equation 7 is that $S_{i,j}$ is a discrete latent variable. This makes designing inference algorithms generally more challenging and potentially unstable for optimisation. A computationally more attractive alternative is the (regularized) horseshoe prior [Carvalho et al., 2009, Piironen and Vehtari, 2017], which serves the same purpose but does not comprise discrete latent variables. In the formulation for a general matrix element $A_{i,j}$, it is defined as

$$A_{i,j} \sim \mathcal{N}\left(0, \frac{C_{i,j} B_{i,j}^2}{C_{i,j} + B_{i,j}^2}\right), \tag{10}$$

$$\tag{11}$$

$$C_{i,j} \sim \mathcal{G}^{-1}(a_0^c, b_0^c), \tag{12}$$

$$B_{i,j} = \Lambda_{i,j} \xi_j \tau, \tag{13}$$

$$\Lambda_{i,j} \sim C^+(0,1), \tag{14}$$

$$\xi_j \sim C^+(0,1), \tag{15}$$

$$\tau \sim C^+(0,1), \tag{16}$$

where $a_0^c = 0.5$ and $b_0^c = 0.5$ are constants with these default values in MOFA-FLEX. The special cases for weights and factor scores priors are again given by $A_{i,j} = W_{d,k}$ for the weights and $A_{i,j} = Z_{n,k}$ for the factor scores. Note, if $C_{i,j} = 0$, we fall back to the unregularized horseshoe prior. The regularized Horseshoe prior in contrast allows for specifying a minimum level of regularization for large coefficients and is used to incorporate prior knowledge in MOFA-FLEX, see Section 1.2.4.

MOFA-FLEX implements both sparsity priors described in this section, but defaults to the horseshoe prior due to its computational advantages.

MOFA-FLEX also implements **Normal** and **Laplace** distributions as weights and factor scores priors, corresponding to L2 and L1 regularisation in a non-Bayesian setting, with the latter inducing variable- or observation-wise sparsity as well. However, they do not account for the multi-view or multi-group structure of the data, and their use corresponds to a simple concatenation of views and groups into a single, unstructured data set.

##### 1.2.2 Spatio-temporal Module

Experiments are often not conducted in complete independence. This is particularly evident in longitudinal studies, where experiments taken close together in time tend to produce more similar results than those taken with larger gaps in time. A similar pattern is observed in spatial omics data, where nearby locations typically show greater similarity than those farther apart. Consequently, the assumption of independent observations used in MOFA/MOFA+ is not suitable in this context. Spatial or temporal information can be provided as additional observation covariates, $\{\mathbf{c}_n\}_{n=1,\dots,N}$ with $\mathbf{c}_n \in \mathbb{R}^P$, where $P$ is the covariate dimensionality (e.g., $P = 1$ for time or $P = 2$ for 2-dimensional space) and $N$ is the number of observations. MEFISTO [Velten et al., 2022] addresses dependent observations by imposing a multivariate prior in the form of a Gaussian process (GP) on each factor in the factor scores matrix. This way, by using an appropriate GP kernel, the factor scores exhibit temporal or spatial smoothness along the covariate. For $G$ observation groups with aligned covariates $\{\mathbf{c}_n^g\}_{n=1,\dots,N_G;g=1,\dots,G}$, MEFISTO additionally implements a multi-group GP kernel that has a covariance component that is shared between groups. To share the same covariate coordinate system between groups, MEFISTO enables the alignment of covariates from multiple groups (in 1D) using the dynamic time warping algorithm [Giorgino, 2009]. MOFA-FLEX adopts the factor prior from MEFISTO, defined as

$$Z_{n,k}^g \sim f_k(\mathbf{c}_n^g) + H_{n,k}^g, \tag{17}$$

$$f_k \sim \mathcal{GP}(0, \kappa_k), \tag{18}$$

$$H^g_{n,k} \sim \mathcal{N}(0, \zeta_k), \tag{19}$$

$$\kappa_k(\mathbf{c}^g_n, \mathbf{c}^{g'}_{n'}) = (1 - \zeta_k)\kappa^C_k(\mathbf{c}^g_n, \mathbf{c}^{g'}_{n'})\kappa^G_k(g, g'), \tag{20}$$

where $\kappa^G_k(g, g')$ is an element of the correlation matrix $\mathbf{K}^G_k$ determined by normalising the group covariance matrix

$$\widetilde{\mathbf{K}}^G_k = \sum_{r=1}^{R} \mathbf{x}^{(k)}_r \mathbf{x}^{(k)T}_r + \sigma^2_k \operatorname{diag}(\mathbf{1}_G) \quad \text{with} \quad \mathbf{x}^{(k)}_r \in \mathbb{R}^G \tag{21}$$

and $\kappa^C_k(\mathbf{c}^g_n, \mathbf{c}^{g'}_{n'})$ is the Radial Basis Function (RBF) kernel used in MEFISTO or the Matérn kernel used in NSF. Both kernels are based on the covariate distance between observations $d = (\mathbf{c}-\mathbf{c}')^{\mathrm{T}}\boldsymbol{\Theta}^{-2}(\mathbf{c}-\mathbf{c}')$, scaled by a lengthscale parameter $\boldsymbol{\Theta}$. The RBF kernel is defined as

$$\kappa^C(\mathbf{c}, \mathbf{c}') = \exp\left(-\frac{1}{2}d\right) \tag{22}$$

and the Matérn kernel is defined as

$$\kappa^C(\mathbf{c}, \mathbf{c}') = \frac{2^{1-\nu}}{\Gamma(\nu)}\left(\sqrt{2\nu}d\right)^{\nu} K_\nu\left(\sqrt{2\nu}d\right), \tag{23}$$

where $\nu \in \{0.5, 1.5, 2.5\}$ is the smoothness parameter and $K_\nu$ is the modified Bessel function. More details about the kernels can be found on the documentation page of GPyTorch: `https://docs.gpytorch.ai/en/v1.13/kernels.html`.

##### 1.2.3 Non-negativity Module

Townes and Engelhardt [2023] introduced Non-negative Spatial Factorisation (NSF), a model that, similarly to MEFIST,O applies Gaussian process priors to the factor scores, but enforces non-negativity of factor scores and weights. This constraint enhances interpretability, particularly for spatial data. However, the exact formulation of NSF is incompatible with MEFISTO's framework. To address this, MOFA-FLEX adheres to MEFISTO's original factor scores prior while offering an optional transformation of factor and weight matrices to non-negative values during inference, using a rectified linear unit (ReLU) [Nair and Hinton, 2010]. Given factor scores $Z_{n,k}$ and weights $W_{n,k}$ from any of the implemented priors, the factorisation with non-negativity constraints is defined as

$$M_{n,d} = \sum_{k=1}^{K} \hat{Z}_{n,k}\hat{W}_{d,k}, \tag{24}$$

$$\hat{Z}_{n,k} = \operatorname{ReLU}(Z_{n,k}), \tag{25}$$

$$\hat{W}_{d,k} = \operatorname{ReLU}(W_{d,k}). \tag{26}$$

Notably, MOFA-FLEX allows independent non-negativity constraints on the factor scores and weights, offering greater flexibility than NSF.

##### 1.2.4 Domain Knowledge Module

Latent variable models with structured sparsity (Section 1.2.1) enable interpretable matrix decomposition. However, assigning biological meaning to factors typically requires additional analysis and can be challenging even for experts. MOFA-FLEX simplifies this process with a domain knowledge module, allowing users to pre-annotate factors with sets of variables (e.g. gene sets from pathway databases) using a modified sparsity prior. This increases the likelihood that factors capture relevant variation, thereby facilitating interpretation.

Prior to model training, a factor is either assigned to a variable set ('informed factor') or remains unassigned ('uninformed factor'). The regularized horseshoe prior (Section 1.2.1) for the weights is then regularized at the factor level, decreasing the prior probability of non-zero weights for variables outside the assigned set. This modification is achieved by element-wise multiplication of the parameter matrix $\mathbf{C}$ (Equation 13) with the squared elements of a scaling matrix $\mathbf{\Omega} \in [0,1]^{D \times K}$, resulting in a new prior for the weights:

$$W_{d,k} \sim \mathcal{N}\left(0, \frac{\widehat{C}_{d,k} B_{d,k}^2}{\widehat{C}_{d,k} + B_{d,k}^2}\right), \tag{27}$$

$$\widehat{C}_{d,k} = C_{d,k} \Omega_{d,k}^2, \tag{28}$$

$$C_{d,k} \sim \mathcal{G}^{-1}(a_0^c, b_0^c), \tag{29}$$

$$B_{d,k} = \Lambda_{d,k} \xi_k \tau, \tag{30}$$

$$\Lambda_{d,k} \sim C^+(0,1), \tag{31}$$

$$\xi_k \sim C^+(0,1), \tag{32}$$

$$\tau \sim C^+(0,1). \tag{33}$$

This way, $\mathbf{\Omega}$ controls the prior scales of individual features and factors. When $\Omega_{d,k} = 1$, the regularized horseshoe prior is recovered, whereas $\Omega_{d,k} \ll 1$ results in a prior scale close to zero, effectively shrinking the weight of feature $d$ for factor $k$ towards zero.

For an uninformed factor $k$, $\Omega_{d,k} = 1$ for all variables $d \in \{1, \dots, D\}$. For an informed factor $k$, $\Omega_{d,k} = 1$ if variable $d$ belongs to the assigned set, and $\Omega_{d,k} = \zeta$ otherwise, where $\zeta \approx 0^+$ serves as a relaxation constant. MOFA-FLEX adopts the default $\zeta = 0.01$ as recommended in the original publication [Qoku and Buettner, 2023]. This relaxation allows features to escape the strong regularization if sufficiently supported by data. Such flexibility is crucial, as predefined variable sets are rarely perfect: Errors in curation and the context-dependent nature of gene programs can both lead to inaccuracies.

###### 1.2.5 Factor Supervision Module

Inferred factor scores often exhibit correlations with external observation covariates, a phenomenon that is both expected and, in many cases, desirable for understanding relationships between covariates and data variables. However, the general matrix factorisation framework (Section 1.1) lacks an explicit mechanism to promote such correlations.

To address this, Capraz et al. [2024] introduced Semi-Supervised Omics Factor Analysis (SOFA), an extension of MOFA that explicitly links individual factors to external covariates. SOFA achieves this by incorporating additional (generalized) linear regression tasks for each covariate, aiming to predict the covariate from the factor scores of a single factor and thereby guiding the factor to model the covariate.

Given the factor scores $Z_{n,k}$ of a single factor $k$ and observation $n$, the linear predictor $\nu_n$ for a given covariate is given by

$$\nu_n = w_0 + w_1 Z_{n,k}, \tag{34}$$

where $w_0$ is an intercept and $w_1$ is a regression coefficient. $w_0$ and $w_1$ are Bayesian latent variables with standard Normal prior distributions:

$$w_i \sim \mathcal{N}(0,1) \quad i \in [0,1]. \tag{35}$$

SOFA supports three types of one-dimensional external covariates $\{c_n\}_{n=1,\dots,N}$:

**Real valued covariates** In this case, $\nu_n$ is the location parameter of a Normal distribution, with the additional scale parameter $\kappa$:

$$c_n \sim \mathcal{N}(\nu_n, \kappa), \tag{36}$$
$$\kappa \sim \mathcal{G}^{-1}(a_0^\kappa, b_0^\kappa), \tag{37}$$

where $a_0^\kappa = e^{-3}$ and $b_0^\kappa = e^{-3}$ are constants.

**Binary covariates** If $c_n \in [0, 1]$, a Bernoulli likelihood with probability parameter $p_n$ given by the logistic function of $\nu_n$ is used:

$$c_n \sim \text{Ber}(p_n), \tag{38}$$
$$p_n = \left(1 + e^{-\nu_n}\right)^{-1}. \tag{39}$$

**Categorical covariates** For a categorical covariate with $R$ categories, $\nu_n^r$ is determined per category with category-specific parameters $w_0^r$ and $w_1^r$, and a Categorical likelihood with probability parameters $p_n^r$ given by the normalized exponential function of $\nu_n^r$ is used:

$$c_n \sim \text{Cat}\left(p_n^1, \ldots, p_n^R\right), \tag{40}$$
$$p_n^r = \left(\sum_r^R e^{\nu_n^r}\right)^{-1} e^{\nu_n^r}. \tag{41}$$

##### 1.2.6 Likelihoods Module

**Normal Likelihood** The default likelihood function in MOFA-FLEX is the Gaussian likelihood $\mathcal{N}\left(\mathbf{M}, \mathbf{\Sigma^2}\right)$ with mean $\mathbf{M} \in \mathbb{R}^{N \times D}$ and diagonal variance $\mathbf{\Sigma^2} \in \mathbb{R}_{\geq 0}^{D \times D}$. While $\mathbf{M}$ is the result of the factorisation, MOFA-FLEX treats $\mathbf{\Sigma_d^2}$ as an additional feature-specific latent variable with an inverse Gamma prior. With this likelihood, the generative model is defined as:

$$Y_{n,d} \sim \mathcal{N}\left(M_{n,d}, \sigma_{d,d}^2\right), \tag{42}$$
$$\sigma_{d,d}^2 \sim \mathcal{G}^{-1}(a_0^\sigma, b_0^\sigma), \tag{43}$$

where $a_0^\sigma = e^{-3}$ and $b_0^\sigma = e^{-3}$ are constants and $M_{n,d}$ is obtained from Equation 4. It is assumed here that $\mathbf{Y}$ is mean-free, otherwise MOFA-FLEX first computes $\mathbf{Y} \leftarrow \mathbf{Y} - \frac{1}{D}\sum_{d=1}^{D} \mathbf{Y_{:,d}}$.

**Bernoulli Likelihood** MOFA-FLEX additionally implements the Bernoulli likelihood for binary input data. With a Bernoulli likelihood, $\mathbf{M}$ is transformed to the unit interval using a logistic function to be a valid probability $\mathbf{P}$. The generative model is defined as:

$$Y_{n,d} \sim \text{Ber}\left(P_{n,d}\right), \tag{44}$$
$$P_{n,d} = 1/(1 + \exp\left(-M_{n,d}\right)), \tag{45}$$

where $M_{n,d}$ is obtained from Equation 4.

**Negative Binomial Likelihood** When using a Negative Binomial likelihood to model count data, MOFA-FLEX transforms $M_{n,d}$ to non-negative values using the ReLU function and scales it by multiplication with the observation mean $\bar{\mathbf{y}} = \sum_n Y_{n,d}/N \in \mathbb{R}^D$. The dispersion parameter $\boldsymbol{\gamma} \in \mathbb{R}_+^D$ is treated as an additional latent variable with a Gamma prior. The generative model is then defined as:

$$
\begin{aligned}
Y_{n,d} &\sim \mathrm{NB}\Big(\widehat{M}_{n,d}, \gamma_d\Big), &(46)\\
\hat{M}_{n,d} &= \mathrm{ReLU}(M_{n,d}) \cdot \bar{y}_d, &(47)\\
\gamma_d^{-1} &\sim \mathcal{G}\big(a_0^\gamma, b_0^\gamma\big), &(48)
\end{aligned}
$$

where $a_0^\gamma = b_0^\gamma = 10^{-10}$ are constants and $M_{n,d}$ is obtained from Equation 4.

#### 1.3 Inference

##### 1.3.1 Stochastic Variational Inference with Pyro

Making downstream inferences from a MOFA-FLEX model requires access to the posterior distributions of the weights $p(\mathbf{W} \mid \mathbf{Y})$ and factor scores $p(\mathbf{Z} \mid \mathbf{Y})$, given the data $\mathbf{Y}$. In the following, we summarise all latent variables in a single random variable $\mathbf{X}$. The posterior distribution can be expressed using Bayes' theorem as:

$$p(\mathbf{X} \mid \mathbf{Y}) = \frac{p(\mathbf{Y} \mid \mathbf{X})p(\mathbf{X})}{p(\mathbf{Y})}, \tag{49}$$

where $p(\mathbf{Y} \mid \mathbf{X})$ is the data likelihood, $p(\mathbf{X})$ is the prior distribution of the latent variables, and $p(\mathbf{Y})$ is the marginal likelihood, also referred to as the model evidence. While this theoretical expression defines the posterior distribution, directly computing $p(\mathbf{Y})$ is typically computationally intractable due to the high-dimensional integrals or summations required.

To overcome this challenge, MOFA-FLEX employs approximate inference via stochastic variational inference (SVI) [Hoffman et al., 2013]. The key idea of SVI is to approximate the true posterior distribution $p(\mathbf{X} \mid \mathbf{Y})$ with a surrogate distribution $q(\mathbf{X})$, referred to as the variational distribution. This approximation is achieved by maximising a computationally feasible objective known as the Evidence Lower Bound (ELBO), which is defined as:

$$\mathrm{ELBO} \coloneqq \mathbb{E}_q[\log p(\mathbf{Y}, \mathbf{X}) - \log q(\mathbf{X})], \tag{50}$$

or equivalently:

$$\mathrm{ELBO} \coloneqq \log p(\mathbf{Y}) - \mathrm{KL}\big(q(\mathbf{X}) \parallel p(\mathbf{X} \mid \mathbf{Y})\big). \tag{51}$$

The ELBO provides a lower bound on the log marginal likelihood, $\log p(\mathbf{Y})$. It contains a Kullback-Leibler (KL) divergence term, a non-negative measure of dissimilarity between the variational distribution and the true posterior. Maximising the ELBO therefore corresponds to simultaneously maximising the log marginal likelihood $\log p(\mathbf{Y})$ and minimising the distance between the variational distribution $q(\mathbf{X})$ and the true posterior $p(\mathbf{X} \mid \mathbf{Y})$.

The ELBO is maximised using stochastic optimisation, which involves (i) parametrising the variational distribution $q(\mathbf{X})$ with a set of learnable parameters $\phi$, (ii) estimating the gradients of the ELBO with respect to $\phi$ using stochastic sampling methods, and (iii) updating the parameters $\phi$ iteratively based on the estimated gradients.

SVI enables scalable inference, making it well-suited for models with large datasets or high-dimensional latent spaces. In MOFA-FLEX, SVI is implemented using the probabilistic programming library Pyro [Bingham et al., 2019], which provides a high-level framework for specifying and optimising probabilistic models. Pyro leverages modern automatic differentiation tools and includes built-in support for variational inference. This streamlined implementation enables efficient experimentation and refinement of the model, facilitating its application to complex datasets.

##### 1.3.2 Inference with Sparse Variational Gaussian Processes in GPyTorch

Gaussian Processes (GPs) are stochastic processes commonly used as non-parametric models for regression and classification. They provide a flexible, probabilistic approach to modelling functions by defining a distribution over functions rather than assuming a fixed parametric form. Formally, a Gaussian Process is a collection of random variables, where any finite subset follows a multivariate Gaussian distribution. A GP is fully characterized by (i) a mean function $m(\mathbf{x})$, which represents the expected value of the function at any input $\mathbf{x}$, and (ii) a covariance function (or kernel) $k(\mathbf{x}, \mathbf{x'})$, which encodes assumptions about the similarity between function values at different points. The choice of kernel governs properties such as smoothness and periodicity. In a Bayesian framework, GPs can be conditioned on observed data points. Given training data, the GP updates its distribution, yielding a posterior that captures both the predicted mean and uncertainty at unobserved points. This makes GPs particularly useful for tasks where uncertainty quantification is important.

Despite their advantages, GPs scale poorly with dataset size $N$, as both computational and memory requirements grow as $\mathcal{O}(N^3)$ and $\mathcal{O}(N^2)$, respectively. This limitation restricts their application to relatively small datasets. Sparse Variational Gaussian Processes (SVGPs) address this scalability issue by introducing a set of $M$ inducing points, where $M \ll N$. These inducing points act as a compact representation of the latent function, reducing computational complexity to $\mathcal{O}(NM^2)$. The SVGP framework employs variational inference to approximate the true posterior distribution of the latent function, optimising a variational evidence lower bound (ELBO). This approach balances model fit and computational efficiency, making GPs applicable to large-scale problems. In MOFA-FLEX, GPs are employed as prior distributions over factor scores. MOFA-FLEX leverages the low-level Pyro interface of the GPyTorch package [Gardner et al., 2018] to seamlessly integrate inference for GPs and the remaining parts of the model.

#### 1.4 Guidance for practitioners

Figure S1 illustrates the modular configuration workflow in MOFA-FLEX. The user configures the model in the following stages.

1. Load data in any compatible format — MuData, AnnData, or standard Python dictionary.
2. Select a Likelihood for each modality, choosing between Normal, Negative Binomial or Bernoulli.
3. Choose a sparsity module for feature selection. We provide a Laplace prior for a simple, parameter-efficient model; a Spike-and-Slab (SNS) prior to emulate classical MOFA-style behaviour; or a (regularised) Horseshoe (HS) prior to incorporate structured domain knowledge from gene programs.
4. Enable the non-negativity module to constrain factor weights to be positive or zero. This is particularly useful when analysing sparse, log-transformed count data, and can enhance interpretability of the latent factors.
5. Activate the domain-knowledge module to infer factors aligned with gene programs. We recommend established resources such as Hallmark and Reactome, though custom gene programs (from GMT files) can be seamlessly integrated.
6. Utilise the Gaussian Process (GP) prior if samples exhibit temporal or spatial structure to learn smooth latent factors over time or tissue space.
7. Account for sample-level covariates such as age, sex, or other clinical variables by introducing guided factors. This supports a controlled and explicit modelling of known sources of variation that may interfere with biologically relevant insight.
8. Define general model parameters, most importantly the number of latent factors. For uninformed models, we recommend training several configurations and comparing the top latent factors by

variance explained. For informed MOFA-FLEX models that ingest gene program annotations to infer biologically meaningful components, 1–3 additional uninformed factors are typically sufficient to absorb residual variation beyond the biological variation explained by the informed programs.

9. Adapt training settings such as the learning rates, number of iterations, mini-batching, and convergence criterion. This step is optionally reserved for advanced users, as MOFA-FLEX already provides meaningful defaults.

10. Explore the trained MOFA-FLEX model utilising the various downstream analysis tools and plotting functionality, such as variance explained, feature importance, factor correlation, factor activity heatmap. These can be used to assess the performance of the model and explore the data. The insights extracted from downstream analyses can be used to update the choice of the model configuration and its parameters.

### 2 Existing Latent Variable Models

The existing landscape of latent variable models is diverse, with various approaches addressing different aspects of data complexity. Table S1 gives an overview of existing methods and their limitations. Notably, MOFA-FLEX stands out by combining multiple sparsity types, pathway priors, non-negativity, spatio-temporal analysis, multi-view and multi-group capabilities, and handling missingness, offering a comprehensive approach to complex data modelling.

### 3 Scalability Benchmarks

We evaluate the GPU-based runtime performance of MOFA-FLEX, ExpiMap, and Spectra across various configurations, as summarized in Table S3 and Figure S2. The experimental setup involves varying the number of annotations (5, 10, 20, and 40) jointly with one other parameter, while keeping all other parameters fixed to the following values: number of samples = 10,000, number of features = 1000, number of factors = 15, and number of views = 1. Note that the number of factors reported is in addition to the number of annotations.

MOFA-FLEX consistently exhibits robust runtime performance with small variability across configurations. All methods demonstrate linear scaling with respect to the number of samples. Notably, Spectra is the only method to exhibit a linear increase in runtime as the number of features grows. The number of views does not significantly impact runtime for any method. However, an increase in the number of factors leads to higher runtimes for ExpiMap and MOFA-FLEX, whereas Spectra maintains a stable runtime under this condition.

Table S3 provides detailed runtime measurements for all configurations. For example, under a configuration with 15 factors, 10,000 samples, and 1,000 features, 1 view and 5 additional annotation, MOFA-FLEX achieves a runtime of approximately 51.71 seconds, outperforming ExpiMap and Spectra, which require 103.17 seconds and 89.21 seconds, respectively. This highlights MOFA-FLEX's efficiency in handling computationally intensive scenarios which occur in many real-world scenarios.

### 4 Additional Results for PBMC Interferon Stimulated Dataset

**Data Preprocessing and Model Training:** To preprocess the data, we selected the 3000 genes with the highest variance across both conditions. For informed gene programs, we chose 65 gene programs from the Reactome and Hallmark databases. We merged gene sets with a Jaccard similarity above 0.8 and filtered those with gene numbers between 40 and 200. We used the `scArches` package (version 0.6.1) for ExpiMap, training it for 1000 epochs with early stopping, a batch size of 1000, and hidden layers of sizes 512 and 256. For Spectra, we utilized the `scSpectra` package (version 0.2.1). Detailed parameters are available in the reproducibility notebooks.

**Comparison to PCA:** To assess the added value of incorporating prior knowledge, we compared the latent representation learned by MOFA-FLEX against a standard PCA baseline. Both methods yielded a clear separation between IFN-stimulated and control cells in UMAP space. However, only MOFA-FLEX achieved this separation through interpretable latent factors directly linked to gene programs via the domain knowledge module. This demonstrates that incorporating structured priors preserves separation performance while providing explicit biological attribution of the underlying variation — a capability that PCA, by construction, cannot offer (Figure S3).

**Gene Program Ranking:** In the PBMC experiment, we demonstrated that MOFA-FLEX's variance explained scores enable meaningful ranking of factors, as evidenced by the high ranking of interferon-named gene programs. Since ExpiMap and Spectra lack a defined measure of variance explained, we investigated whether the L2-Norm of the factor loadings could serve as an alternative for guiding factor selection. Although interferon-named gene programs are among the top 20 gene programs for ExpiMap and Spectra, they do not occupy the highest ranks, as shown in Figure S4.

**Choice of Number of Factors:** To address the challenge of selecting the appropriate number of factors in factor analysis, we examined ability of the L2-Norm and Variances Explained used by the three methods in Figure S5. Since there is no notion of variance explained in ExpiMap and Spectra, we selected the L2-Norm as an alternative. We created synthetic data based on $K = 10$ (informed) latent factors and trained all models with $K = 15$ factors. ExpiMap and Spectra employ the L2-Norm of factor loadings, with values that do not allow to observe a clear cut-off after 10 factors. In contrast, MOFA-FLEX's variance explained illustrates a ranking of factors showing a clear cut-off after 10 factors (dropping to nearly 0) and thereby allowing to guide the parameter choice.

**Synthetic Experiments** For the synthetic data experiments, we utilised 65 gene programs from the Hallmark and Reactome databases. We generated 10000 synthetic data points by sampling $z_{ng} \sim \mathcal{N}(0, 1)$, normalising the values, and introducing varying levels of false positives and false negatives to the weights $W$ (representing the ground-truth Hallmark and Reactome gene programs). We then computed $Y = WZ + \epsilon$, where $\epsilon_{gn} \sim \mathcal{N}(0, 1)$. This data was provided to the three models, and we evaluated their ability to reconstruct the ground-truth $W$ using AUPRC and their reconstruction accuracy for the input data $Y$.

**Inference of Novel Genes in IFN-signatures** By integrating prior information with data-driven evidence, MOFA-FLEX extends canonical gene program signatures with novel, biologically coherent gene candidates. In addition to the gene programs reported in the main section, we report further gene programs with the highest variance explained scores in Figure S6. For example, MOFA-FLEX refined "Interferon Alpha Beta Signaling" gene set from Reactome with the (1) GBP1, GBP4 and GBP5 genes. GBP1 mediates antiviral effects and therefore plays a key role in the immune response, and its expression is upregulated due to the stimulation [Li et al., 2024]. GBP4 is also an interferon-inducible gene and acts as a negative regulator of type I interferon responses by interacting with IRF7 and inhibiting its ability to respond, thereby modulating the magnitude of the antiviral response. Overexpression of GBP4 dampens IFN-I signaling, while its knockdown enhances it [Hu et al., 2011]. GBP5 is upregulated in PBMCs following IFN-$\beta$ stimulation and contributes to antiviral defense by activating interferon signaling gene programs and promoting the expression of pro-inflammatory cytokines and chemokines [Feng et al., 2017]. As another example, CCL8 (chemokine ligand 8) is induced in PBMCs by interferon stimulation, including IFN-$\beta$ and IFN-$\gamma$ (see the contextualised "Interferon Gamma Response" gene program from Hallmark). It is part of the chemokine response that mediates immune cell recruitment during antiviral responses, i.e. recruiting monocytes, macrophages, and other immune cells to sites of inflammation. Moreover, considering the "Interferon Alpha Response" Hallmark gene program, we consider the contextualisation of e.g., IFI6 (Interferon Alpha Inducible Protein 6) an interferon-stimulated gene, which is induced by type I interferons, including interferon beta. IFI6 acts as a negative regulator of innate immune responses by modulating RIG-I activation and suppressing excessive interferon and inflammatory cytokine expression, thereby contributing to the feedback regulation of the interferon alpha response gene program [Villamayor et al., 2023, Sajid et al., 2021].

### 5 Handling Technical Variation and Noisy Priors in the Multiomic CITE-seq Data

In this section, we apply MOFA-FLEX with alternative training settings to highlight two key aspects of our method. First, in the presence of biological prior information, MOFA-FLEX effectively redistributes sources of non-biological or technical variation, such as batch effects, onto the designated uninformed latent factors. In contrast, a standard uninformed MOFA-FLEX model disperses technical variance across multiple factors, rather than confining it to the top few factors that are suitable for differentiating between the batches (Figure S7). By integrating gene set annotations a priori, MOFA-FLEX extends canonical cell-type signatures with novel, biologically coherent gene candidates. In addition to the gene programs reported in the main section, we report further gene programs with the highest variance explained scores in Figure S8.

Second, MOFA-FLEX demonstrates robustness to varying levels of noise in the prior information, refining noisy annotations to more accurately reflect the underlying gene program composition. To evaluate this, we applied MOFA-FLEX to the CITE-seq dataset of immune cells from murine spleen and lymph nodes, incorporating prior annotations derived from mouse cell type signatures specific to ageing processes [tab, 2020], particularly within the spleen. Consistent with the main analysis, we informed MOFA-FLEX with Hallmark annotations from the mouse MSigDB [Subramanian et al., 2005, Liberzon et al., 2015, Castanza et al., 2023] and cell type signatures from the tissue-specific mouse cell atlas (Tabula Muris) [tab, 2020], resulting in 62 distinct gene sets with a median size of 67 genes, closely mirroring the original training conditions. The efficacy of the informed MOFA-FLEX models is also evident in the results obtained by the single-cell integration benchmarking (scIB) [Luecken et al., 2022] as shown in the Figure S9. In particular, the informed MOFA-FLEX models achieve a better separation between the non-biological factors and biologically informed factors which preserve relevant biological information. MOFA-FLEX was trained using default parameters, including three additional uninformed factors to capture variance beyond gene program-associated signals. The comparison results between models are presented in Figure S10, highlighting MOFA-FLEX's ability to refine noisy priors and enhance biological interpretability.

### 6 Overcoming Low Gene Coverage in Spatial Assays via Joint Modelling of Spatial and Single-cell Transcriptomics

The advantage of integrating gene sets with spatial information a priori is two-fold; First, the inference of the latent factors is guided by relevant domain knowledge from suitable biological processes. Second, the presence of cell coordinates render the inferred factors spatially aware, thereby localizing their activity in meaningful areas across the tissue. Here, we highlight how MOFA-FLEX leverages multimodal integration to overcome the limitations of low gene coverage in spatial assays. While the Xenium assay provides precise spatial localization of gene expression, its limited panel of ∼300 genes constrains the depth of gene program analysis. To address this, we incorporate high-coverage Chromium single-cell transcriptomics (3,000 genes), effectively borrowing genes to enrich the prior information available for jointly modelling both groups. This approach significantly enhances gene program detection, as evidenced by Principal Component Gene Set Enrichment (PCGSE) [Frost et al., 2015] analysis: whereas applying PCGSE to Xenium alone fails to yield significant factors, the joint modelling of Xenium and Chromium renders virtually all factors highly significant. A comprehensive comparison of the inferred gene programs is provided in Figure S11, where the majority of the loadings when mapping the prior annotations onto the Xenium (low gene coverage) are inferred.

Beyond gene program detection, this integration also improves gene program-based classification performance. Notably, for distinguishing cancerous from non-cancerous cells, the latent factor that was initially informed by the Estrogen Response Early (ERE) gene program maintains a high AUROC of 0.97 in both the Xenium-only and Xenium + Chromium models, indicating that the available Xenium genes might be sufficient for this classification task. However, for the latent factor informed by the Apical Junction (AJ) gene program, which differentiates invasive tumor from non-invasive tumor (DCIS 1 and DCIS 2), the AUROC improves substantially from 0.83 in the Xenium-only model to 0.93 in the multi-group model. This improvement is driven by the expanded gene set: while Xenium alone includes only 6 genes from the AJ gene program, integrating Chromium increases coverage to 66 genes, enriching the prior information and enhancing MOFA-FLEX's ability to capture biologically meaningful spatial patterns. To further explore the dataset, we focus on the multi-group application, and examine the biological gene programs for markers that are potentially relevant in the diagnosis and prognosis of breast cancer. Among the top annotated genes in the Estrogen Response Early (ERE) gene program, we observe several genes with significant implications in breast cancer diagnosis, prognosis, and survival. TFF1 [Yi et al., 2020], GATA3 [Takaku et al., 2015], and FOXA1 [Metovic et al., 2022] are classic estrogen-regulated markers associated with luminal breast cancer subtypes. TFF1 is upregulated in breast cancer tissue and can predict the prognosis of patients with breast cancer. Forkhead box A1 protein (FOXA1) is essential for ER chromatin binding, and plays a crucial role in the development and progression of breast cancer. In addition, FASN [Vanauberg et al., 2023], CCND1 [Valla et al., 2022], and ERBB2 [Revillion et al., 1998, Tan and Yu, 2007] are associated with cell proliferation, metabolic rewiring, and poor prognosis. The fatty acid synthase (FASN), responsible for the increased production of fatty acids, is overexpressed in breast cancer by promoting carcinogenesis, cell proliferation and breast cancer metastasis. The ERBB2 receptor, also referred to as HER2, correlates strongly with the aggressiveness of the cancer and is strongly associated with poor prognosis in breast cancer patients. On the other hand, ESRP2 [He et al., 2024] and CLDN7 [Fan et al., 2024] contribute to epithelial cell maintenance, while their dysregulation is linked to metastasis. The inhibition of ESRP2, in particular, is directly linked to a decrease in cyclinD1 protein expression, encoded by the CCND1 gene. Furthermore, overexpression of ESRP2 is correlated with poor prognosis in breast cancer patients. Next, the epithelial cell adhesion molecule (EpCAM) [Osta et al., 2004] is commonly expressed in breast cancer, and has been linked to poor prognosis of patients with breast cancer. Moreover, the refined Apical Junction (AJ) gene program, enriched in ductal carcinoma in situ (DCIS) compared to invasive tumour cells, reinforces the critical role of cell-cell adhesion, apical polarity, and cytoskeletal integrity in maintaining epithelial homeostasis and restricting tumour migration and progression. Key genes such as CEACAM6 [Lewis-Wambi et al., 2008] and TACSTD2 [Vidula et al., 2022] are prominently expressed in DCIS, promoting epithelial polarity and adhesion, while their downregulation in invasive tumor cells facilitates epithelial-to-mesenchymal transition (EMT), enabling motility, invasion and subsequent metastasis. Tight junction proteins like claudins (CLDN4) and adhesion molecules such as JUP [Holen et al., 2012] stabilize apical junction complexes in DCIS, ensuring the compartmentalization of epithelial layers. However, their loss was shown to lower cell–cell contact, thereby promoting detachment and interaction with the surrounding stromal environment. Actinins, including ACTN1 [Kovac et al., 2018], ACTN4 [Hsu and Kao, 2013], and ACTG1, are cytoskeletal proteins that have been shown to regulate cytokinesis, cell adhesion, tumorigenesis and invasive potential in breast cancer, with potential for being ideal drug targets for future therapeutic development. Furthermore, the diminished expression of classical epithelial markers, such as CDH1 [Lombaerts et al., 2006], in invasive tumor cells emphasizes the breakdown of epithelial integrity during EMT. These findings underscore the critical role of apical junction components in distinguishing DCIS from invasive phenotypes and suggest their potential as biomarkers or therapeutic targets in breast cancer. By integrating complementary data sources, MOFA-FLEX not only improves gene program resolution but also enables a more robust and interpretable mapping of transcriptome-wide activity across spatial tissue architecture, providing deeper insights into tumor biology and disease progression.

### 7 Reproduction of MOFA, MEFISTO and NSF Results

MOFA-FLEX provides modules to replicate several published matrix factorisation models. To demonstrate that MOFA-FLEX reliably reproduces results from the original publications, we selected data sets used in MOFA, MEFISTO, and NSF and applied MOFA-FLEX with a module configuration roughly matching the original model.

#### 7.1 MOFA

We ran MOFA-FLEX and MOFA on the chronic lymphocytic leukaemia (CLL) dataset [Dietrich et al., 2018] featured in the MOFA publication. This dataset comprises transcriptome, DNA methylation, somatic mutation, and *ex vivo* drug response measurements from 200 samples with a varying number of missing data.

We trained the MOFA model using the `mofapy2` package (version 0.7.2) with 10 factors, a convergence mode 'medium', and otherwise default parameters. For MOFA-FLEX, we trained 10 models with different initialisation seeds, each configured with 10 factors, a Horseshoe prior for the weights, a Gaussian prior for the factor scores, a Gaussian data likelihood for all views, a learning rate of 0.05, and default parameters otherwise.

For each trained MOFA-FLEX model, we aligned factors and their sign with those of MOFA to maximise the correlation of factor scores, using the `linear_sum_assignment` function in SciPy applied to the absolute values of the cross-correlation matrix. We then computed the Pearson correlation coefficient of weights and factor scores per factor (Figure S12a), showing generally good correlation. We further validated that MOFA-FLEX captures the strong signals in the weights of the mutations view observed in the MOFA publication. Specifically, del11q22.3 and IGHV were identified as key features in the first factor, while del13q14_any and Trisomy 12 were prominent in the second factor (Figure S12b). The distribution of explained variance across factors and views was similar between MOFA-FLEX and MOFA (Figure S12c). Additionally, MOFA-FLEX successfully reproduced the separation of samples by IGHV and Trisomy 12 base on factors 1 and 2, yielding results closely aligned with those from MOFA (Figure S12).

#### 7.2 MEFISTO

We ran MOFA-FLEX and MEFISTO on the 10X Visium Spatial Transcriptomics dataset of the anterior mouse brain [Stuart et al., 2019], as analysed in the MEFISTO publication. We trained a MEFISTO model using the `mofapy2` package (version 0.7.2) with 4 factors, sparse Gaussian processes with 1000 inducing points, `start_opt=10`, `opt_freq=30`, and otherwise default parameters. For MOFA-FLEX, we trained 10 models with different initialisation seeds, each configured with 4 factors, a Horseshoe prior for the weights, a Gaussian process prior with RBF kernel and 1000 inducing points for the factor scores, a Gaussian data likelihood, a learning rate of $lr = 0.05$, and default parameters otherwise.

We aligned the factors in the same way as for MOFA before and computed the Pearson correlation coefficient of weights and factor scores per factor (Figure S13a), showing high correlation and low variance across MOFA-FLEX initialisations. We visualized the weights individually for different factors, showing strong correlations between MOFA-FLEX and MEFISTO for all four factors (Figure S13b).

Finally, we visualised factor scores with their spatial coordinates, revealing highly concordant spatial patterns between the two models (Figure S13c).

###### 7.2.1 Inferring Biologically Informed Latent Factors

In addition to the uninformed MOFA-FLEX application, we trained a MOFA-FLEX model on the mouse brain dataset with prior gene set annotations from PanglaoDB [Franzén et al., 2019]. We filtered for the subset of signatures pertaining to mice brain cell types, resulting in 24 cell type signatures in total. As shown in Figure S14, the informed MOFA-FLEX model revealed spatially aware gene program activity across the anterior mouse brain. Among others, we observed different areas enriched with neuroendocrine cells, Purkinje neurons, interneurons and choroid plexus cells. In particular, the choroid plexus cells are driven by the TTR (plasma transthyretin) marker, which is highly abundant in the choroid plexus [Herbert et al., 1986] (Fig. S14b).

##### 7.3 NSF

We ran MOFA-FLEX on the Slide-seq V2 data of the mouse hippocampus that was used in the NSF publication. It comprises 365,360 spatial observations and 2000 genes (after the same preprocessing as in the NSF publication). The MOFA-FLEX model had 6 factors, a Horseshoe prior for the weights, a Gaussian process prior for the factor scores with Matern kernel and 500 inducing points, a Gamma-Poisson data likelihood, nonnegativity constraints on weights and factor scores, mini-batch training with a batch size of 1000 observations, and a learning rate of $lr = 5e - 3$.

The obtained factors strongly resemble the shown factors in [Townes and Engelhardt, 2023] (Figure S15a) and we conclude that in a similar configuration (notably not exactly the same as NSF because of incompatibilities of the MEFISTO and NSF factor scores priors), MOFA-FLEX can reproduce the results from the original publication.

#### 8 Code Example

MOFA-FLEX was designed to allow the rapid prototyping of custom matrix factorization models, enabling researchers to efficiently define and iterate on new models in the field of computational biology. Moreover, the MOFA-FLEX framework allows its users to easily share their models with other researchers via our codebase. An important feature therefore is the intuitive interface to define the models, including the integration of covariates, such as spatial or temporal dependencies between samples. Hence, we provide an interface that allows (i) importing published models, such as MOFA or MEFISTO, (ii) defining models with little effort by relying on our set of predefined parameters and (iii) customizing each detail of the model for expert users.
In the following code example, a MOFA-FLEX model is trained on 2 data groups with two views each (RNA and protein). The data does not contain any covariates, and the genes in the RNA view are informed with gene sets from a `.gmt` file.

```python
from mofaflex import MOFAFLEX, ModelOptions, TrainingOptions, FeatureSets

# prepare the data, which is a nested dictionary of AnnData objects
data = {
    "group_1" : {"rna" : adata_rna_g1, "prot" : adata_prot_g1},
    "group_2" : {"rna" : adata_rna_g2, "prot" : adata_prot_g2},
}

# load and filter gene sets by overlap with available genes
gene_set_collection = FeatureSets.from_gmt(...).filter(
    data["group_1"]["view_rna"].var_names,
    min_fraction=0.05,
    min_count=10,
)

# create gene set masks for the RNA view in each group
for group in ["group_1", "group_2"]:
    data[group]["rna"].varm["gene_set_mask"] = gene_set_collection.to_mask(
        data[group]["rna"].var_names.tolist()
    ).T

# instantiate and train a MOFA-FLEX model
model = MOFAFLEX(
    data,
    ModelOptions(
        n_factors=10,
        weight_prior={"view_rna" : "Horseshoe", "view_prot" : "Laplace"},
        factor_prior="Normal",
        likelihoods={"view_rna" : "GammaPoisson", "view_prot" : "Normal"},
        nonnegative_weights=False,
        annotations_varm_key={"view_rna" : "gene_set_mask"},
    ),
    TrainingOptions(
        device="cuda:0",
        lr=5e-3,
    )
)
```

Oscar Franzén, Li-Ming Gan, and Johan LM Björkegren. Panglaodb: a web server for exploration of mouse and human single-cell rna sequencing data. *Database*, 2019:baz046, 2019.

H Robert Frost, Zhigang Li, and Jason H Moore. Principal component gene set enrichment (pcgse). *BioData mining*, 8(1):25, 2015.

Jacob Gardner, Geoff Pleiss, Kilian Q Weinberger, David Bindel, and Andrew G Wilson. Gpytorch: Blackbox matrix-matrix gaussian process inference with gpu acceleration. *Advances in neural information processing systems*, 31, 2018.

Adam Gayoso, Zoë Steier, Romain Lopez, Jeffrey Regier, Kristopher L Nazor, Aaron Streets, and Nir Yosef. Joint probabilistic modeling of single-cell multi-omic data with totalvi. *Nature methods*, 18 (3):272–282, 2021.

Toni Giorgino. Computing and visualizing dynamic time warping alignments in r: the dtw package. *Journal of statistical Software*, 31:1–24, 2009.

Xiaoping Han, Renying Wang, Yincong Zhou, Lijiang Fei, Huiyu Sun, Shujing Lai, Assieh Saadatpour, Ziming Zhou, Haide Chen, Fang Ye, et al. Mapping the mouse cell atlas by microwell-seq. *Cell*, 172(5):1091–1107, 2018.

Caiping He, Yuting Chen, Ximin Zhang, Huancun Feng, Yuzhen Rao, Tangyang Ji, and Wenya Wang. Down-regulation of esrp2 inhibits breast cancer cell proliferation via inhibiting cyclind1. *Scientific Reports*, 14(1):28475, 2024.

Joseph Herbert, Josiah N Wilcox, Kim-Thu C Pham, Robert T Fremeau Jr, Massimo Zeviani, Andrew Dwork, Dianne R Soprano, Adina Makover, DeWitt S Goodman, Earl A Zimmerman, et al. Transthyretin: A choroid plexus-specific transport protein in human brain: The 1986 s. weir mitchell award. *Neurology*, 36(7):900–900, 1986.

Matthew D Hoffman, David M Blei, Chong Wang, and John Paisley. Stochastic variational inference. *Journal of Machine Learning Research*, 2013.

Ingunn Holen, Jacob Whitworth, Faith Nutter, Alyson Evans, Hannah K Brown, Diane V Lefley, Ivana Barbaric, Mark Jones, and Penelope D Ottewell. Loss of plakoglobin promotes decreased cell-cell contact, increased invasion, and breast cancer cell dissemination in vivo. *Breast Cancer Research*, 14:1–14, 2012.

Kuo-Sheng Hsu and Hung-Ying Kao. Alpha-actinin 4 and tumorigenesis of breast cancer. *Vitamins & Hormones*, 93:323–351, 2013.

Yu Hu, Jie Wang, Bo Yang, Nuoyan Zheng, Meiling Qin, Yongyong Ji, Guomei Lin, Lin Tian, Xiaodong Wu, Li Wu, et al. Guanylate binding protein 4 negatively regulates virus-induced type i ifn and antiviral response by targeting ifn regulatory factor 7. *The Journal of Immunology*, 187(12): 6456–6462, 2011.

Amanda Janesick, Robert Shelansky, Andrew D Gottscho, Florian Wagner, Stephen R Williams, Morgane Rouault, Ghezal Beliakoff, Carolyn A Morrison, Michelli F Oliveira, Jordan T Sicherman, et al. High resolution mapping of the tumor microenvironment using integrated single-cell, spatial and in situ analysis. *Nature communications*, 14(1):8353, 2023.

Hyun Min Kang, Meena Subramaniam, Sasha Targ, Michelle Nguyen, Lenka Maliskova, Elizabeth McCarthy, Eunice Wan, Simon Wong, Lauren Byrnes, Cristina M Lanata, et al. Multiplexed droplet single-cell rna-sequencing using natural genetic variation. *Nature biotechnology*, 36(1):89–94, 2018.

Bianca Kovac, Tomi P Mäkelä, and Tea Vallenius. Increased $\alpha$-actinin-1 destabilizes e-cadherin-based adhesions and associates with poor prognosis in basal-like breast cancer. *PloS one*, 13(5):e0196986, 2018.

Russell Z Kunes, Thomas Walle, Max Land, Tal Nawy, and Dana Pe'er. Supervised discovery of interpretable gene programs from single-cell data. *Nature Biotechnology*, 42(7):1084–1095, 2024.

Joan S Lewis-Wambi, Heather E Cunliffe, Helen R Kim, Amanda L Willis, and V Craig Jordan. Overexpression of ceacam6 promotes migration and invasion of oestrogen-deprived breast cancer cells. *European journal of cancer*, 44(12):1770–1779, 2008.

Yadi Li, Haiying Luo, Xiaoxia Hu, Jiaojiao Gong, Guili Tan, Huating Luo, Rui Wang, Hao Pang,

Renjie Yu, and Bo Qin. Guanylate-binding protein 1 (gbp1) enhances ifn-$\alpha$ mediated antiviral activity against hepatitis b virus infection. *Polish Journal of Microbiology*, 73(2):217, 2024.

Arthur Liberzon, Chet Birger, Helga Thorvaldsdóttir, Mahmoud Ghandi, Jill P Mesirov, and Pablo Tamayo. The molecular signatures database hallmark gene set collection. *Cell systems*, 1(6):417–425, 2015.

M Lombaerts, T Van Wezel, K Philippo, JWF Dierssen, RME Zimmerman, J Oosting, R Van Eijk, PH Eilers, B van De Water, CJ Cornelisse, et al. E-cadherin transcriptional downregulation by promoter methylation but not mutation is related to epithelial-to-mesenchymal transition in breast cancer cell lines. *British journal of cancer*, 94(5):661–671, 2006.

Mohammad Lotfollahi, Sergei Rybakov, Karin Hrovatin, Soroor Hediyeh-Zadeh, Carlos Talavera-López, Alexander V Misharin, and Fabian J Theis. Biologically informed deep learning to query gene programs in single-cell atlases. *Nature Cell Biology*, 25(2):337–350, 2023.

Malte D Luecken, Maren Büttner, Kridsadakorn Chaichoompu, Anna Danese, Marta Interlandi, Michaela F Müller, Daniel C Strobl, Luke Zappia, Martin Dugas, Maria Colomé-Tatché, et al. Benchmarking atlas-level data integration in single-cell genomics. *Nature methods*, 19(1):41–50, 2022.

David JC MacKay. Bayesian methods for backpropagation networks. In *Models of neural networks III: association, generalization, and representation*, pages 211–254. Springer, 1996.

Jasna Metovic, Fulvio Borella, Marta D'Alonzo, Nicoletta Biglia, Luca Mangherini, Cristian Tampieri, Luca Bertero, Paola Cassoni, and Isabella Castellano. Foxa1 in breast cancer: a luminal marker with promising prognostic and predictive impact. *Cancers*, 14(19):4699, 2022.

Toby J Mitchell and John J Beauchamp. Bayesian variable selection in linear regression. *Journal of the american statistical association*, 83(404):1023–1032, 1988.

Vinod Nair and Geoffrey E Hinton. Rectified linear units improve restricted boltzmann machines. In *Proceedings of the 27th international conference on machine learning (ICML-10)*, pages 807–814, 2010.

Walid A Osta, Yian Chen, Kaidi Mikhitarian, Michael Mitas, Mohamed Salem, Yusuf A Hannun, David J Cole, and William E Gillanders. Epcam is overexpressed in breast cancer and is a potential target for breast cancer gene therapy. *Cancer research*, 64(16):5818–5824, 2004.

Emma Pierson and Christopher Yau. Zifa: Dimensionality reduction for zero-inflated single-cell gene expression analysis. *Genome biology*, 16:1–10, 2015.

Juho Piironen and Aki Vehtari. Sparsity information and regularization in the horseshoe and other shrinkage priors. *Electronic Journal of Statistics*, 11(2):5018 – 5051, 2017. doi: 10.1214/17-EJS1337SI. URL `https://doi.org/10.1214/17-EJS1337SI`.

Arber Qoku and Florian Buettner. Encoding domain knowledge in multi-view latent variable models: a bayesian approach with structured sparsity. In *International Conference on Artificial Intelligence and Statistics*, pages 11545–11562. PMLR, 2023.

F Revillion, J Bonneterre, and JP Peyrat. Erbb2 oncogene in human breast cancer and its clinical significance. *European Journal of Cancer*, 34(6):791–808, 1998.

Davide Risso, Fanny Perraudeau, Svetlana Gribkova, Sandrine Dudoit, and Jean-Philippe Vert. A general and flexible method for signal extraction from single-cell rna-seq data. *Nature communications*, 9(1):284, 2018.

Muhammad Sajid, Hafiz Ullah, Kun Yan, Miao He, Jiangpeng Feng, Muhammad Adnan Shereen, Ruidong Hao, Qiaohong Li, Deyin Guo, Yu Chen, et al. The functional and antiviral activity of interferon alpha-inducible ifi6 against hepatitis b virus replication and gene expression. *Frontiers in immunology*, 12:634937, 2021.

Rahul Satija, Paul Hoffman, and Andrew Butler. *SeuratData: Install and Manage Seurat Datasets*, 2025. URL `https://github.com/satijalab/seurat-data`. R package version 0.2.2.9002.

Tim Stuart, Andrew Butler, Paul Hoffman, Christoph Hafemeister, Efthymia Papalexi, William M Mauck, Yuhan Hao, Marlon Stoeckius, Peter Smibert, and Rahul Satija. Comprehensive integration of single-cell data. *cell*, 177(7):1888–1902, 2019.

Aravind Subramanian, Pablo Tamayo, Vamsi K Mootha, Sayan Mukherjee, Benjamin L Ebert, Michael A Gillette, Amanda Paulovich, Scott L Pomeroy, Todd R Golub, Eric S Lander, et al. Gene set enrichment analysis: a knowledge-based approach for interpreting genome-wide expression profiles. *Proceedings of the National Academy of Sciences*, 102(43):15545–15550, 2005.

Motoki Takaku, Sara A Grimm, and Paul A Wade. Gata3 in breast cancer: tumor suppressor or oncogene? *Gene expression*, 16(4):163, 2015.

Ming Tan and Dihua Yu. Molecular mechanisms of erbb2-mediated breast cancer chemoresistance. *Breast Cancer Chemosensitivity*, pages 119–129, 2007.

F William Townes and Barbara E Engelhardt. Nonnegative spatial factorization applied to spatial genomics. *Nature methods*, 20(2):229–238, 2023.

F William Townes, Stephanie C Hicks, Martin J Aryee, and Rafael A Irizarry. Feature selection and dimension reduction for single-cell rna-seq based on a multinomial model. *Genome biology*, 20:1–16, 2019.

Marit Valla, Elise Klæstad, Borgny Ytterhus, and Anna M Bofin. Ccnd1 amplification in breast cancer-associations with proliferation, histopathological grade, molecular subtype and prognosis. *Journal of mammary gland biology and neoplasia*, 27(1):67–77, 2022.

Dimitri Vanauberg, Céline Schulz, and Tony Lefebvre. Involvement of the pro-oncogenic enzyme fatty acid synthase in the hallmarks of cancer: a promising target in anti-cancer therapies. *Oncogenesis*, 12(1):16, 2023.

Britta Velten, Jana M Braunger, Ricard Argelaguet, Damien Arnol, Jakob Wirbel, Danila Bredikhin, Georg Zeller, and Oliver Stegle. Identifying temporal and spatial patterns of variation from multimodal data using mefisto. *Nature methods*, 19(2):179–186, 2022.

Neelima Vidula, Christina Yau, and Hope Rugo. Trophoblast cell surface antigen 2 gene (tacstd2) expression in primary breast cancer. *Breast Cancer Research and Treatment*, 194(3):569–575, 2022.

Laura Villamayor, Vanessa Rivero, Darío López-García, David J Topham, Luis Martínez-Sobrido, Aitor Nogales, and Marta L DeDiego. Interferon alpha inducible protein 6 is a negative regulator of innate immune responses by modulating rig-i activation. *Frontiers in immunology*, 14:1105309, 2023.

Florin C Walter, Oliver Stegle, and Britta Velten. Fishfactor: a probabilistic factor model for spatial transcriptomics data with subcellular resolution. *Bioinformatics*, 39(5):btad183, 2023.

Jie Yi, Liwen Ren, Dandan Li, Jie Wu, Wan Li, Guanhua Du, and Jinhua Wang. Trefoil factor 1 (tff1) is a potential prognostic biomarker with functional significance in breast cancers. *Biomedicine & Pharmacotherapy*, 124:109827, 2020.
